# Multi-scale Adaptive Differential Abundance Analysis in Microbial Compositional Data

**DOI:** 10.1101/2021.11.02.466987

**Authors:** Shulei Wang

## Abstract

Differential abundance analysis is an essential and commonly used tool to characterize the difference between microbial communities. However, identifying differentially abundant microbes remains a challenging problem because the observed microbiome data is inherently compositional, excessive sparse, and distorted by experimental bias. Besides these major challenges, the results of differential abundance analysis also depend largely on the choice of analysis unit, adding another practical complexity to this already complicated problem. In this work, we introduce a new differential abundance test called the MsRDB test, which embeds the sequences into a metric space and integrates a multi-scale adaptive strategy for utilizing spatial structure to identify differentially abundant microbes. Compared with existing methods, the MsRDB test can detect differentially abundant microbes at the finest resolution offered by data and provide adequate detection power while being robust to zero counts, compositional effect, and experimental bias in the microbial compositional data set. Applications to both simulated and real microbial compositional data sets demonstrate the usefulness of the MsRDB test.

## 1 Introduction

Differential abundance analysis is a commonly used tool to decipher the difference between microbial communities and identify dysbiotic microbes. The primary goal of differential abundance analysis is to detect a set of microbes associated with experimental conditions based on observed count data generated from high-throughput sequencing technologies. Although popular, differential abundance analysis in microbiome data remains a challenging statistical problem as the observed count data is inherently compositional (Li, 2015; Gloor et al., 2017; Vandeputte et al., 2017; Morton et al., 2019), excessive sparse (Brill et al., 2022), and distorted by experimental bias (McLaren et al., 2019) (Figure 1). To address these challenges, a number of studies have proposed differential abundance analysis procedures for microbiome data (Paulson et al., 2013; Fernandes et al., 2013; Mandal et al., 2015; Shi et al., 2016; Tang et al., 2017; Xiao et al., 2017; Chen et al., 2018; Morton et al., 2019; Brill et al., 2022; Lin and Peddada, 2020a; Martin et al., 2020; Zhou et al., 2021; Wang, 2023; Zhou et al., 2022). See a comprehensive review of differential abundance analysis in Weiss et al. (2017) or Lin and Peddada (2020b).

**Figure 1:**
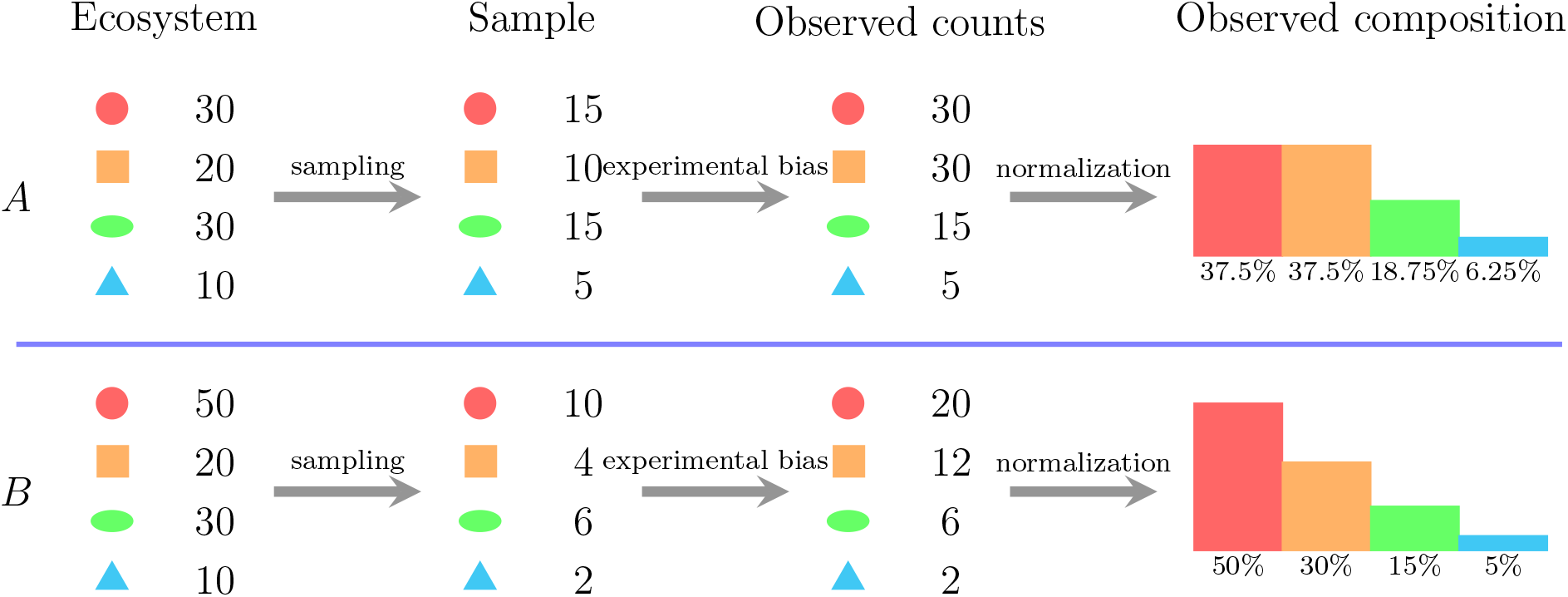
The limitations inherent in the observed microbial compositional data set. This figure shows that the sampling process and experimental bias in sequencing workflow create a systemic distinction between absolute abundance in the ecosystem and our observed count data. The observed composition after normalization can also be significantly different from the relative abundance in the original ecosystem owing to the experimental bias. The measurement efficiency in this example is (2, 3, 1, 1).

Before applying these existing methods, one must choose an appropriate analysis unit for differential abundance analysis. The reads in 16S rRNA sequencing data are clustered into operational taxonomic units (OTUs) (Hamady and Knight, 2009) or denoised as amplicon sequence variants (ASVs) (Callahan et al., 2016; Amir et al., 2017). Using OTUs or ASVs as the analysis unit, we can detect the differentially abundant microbes at the finest resolution offered by 16S rRNA sequencing data (Figure 3A). However, such a strategy sometimes can suffer a loss of detection power as the sparse counts in each OTU or ASV are not informative enough for an effective comparison (Huang et al., 2021). To increase power, one commonly used strategy is to aggregate similar OTUs or ASVs as taxa, such as genera, according to their assigned/mapped taxonomy (Figure 3B). Treating taxa as analysis units can provide more power to detect differentially abundant microbes but at a coarser resolution. Choosing a taxonomy rank in the taxon-wise analysis also needs facing a tradeoff between resolution and sequence utilization. On the one hand, a lower taxonomy rank can provide finer resolution but requires throwing out many unassigned sequences due to the current coarse taxonomic classification. On the other hand, a higher taxonomy rank can keep and utilize more sequences but loses resolution to capture the heterogeneous signal. Besides aggregating based on assigned taxonomy, another strategy to combine the OTUs or ASVs is to incorporate the hierarchical structure of the phylogenetic tree (Yekutieli, 2008; Tang et al., 2017; Xiao et al., 2017; Heller et al., 2018; Bichat et al., 2020; Li et al., 2020; Huang et al., 2021). Such a strategy allows detection of the differential abundant microbes at different resolutions and improves the detection power compared with OTU/ASV-wise analysis. However, this strategy usually requires accurate evolution information of the phylogenetic tree, and it is still unknown how the error in phylogenetic tree construction affects the downstream differential abundance analysis. It is natural to wonder if there is a way to detect differentially abundant microbes at OTU or ASV resolution robustly while maintaining enough power and using all available sequences. Motivated by this need, we introduce an alternative way to conduct differential abundance analysis in this paper.

Here, we propose the MsRDB test (Multiscale Adaptive Robust Differential Abundance Test), a multi-scale method for differential abundance analysis in microbial compositional data. Instead of independent units, the MsRDB test embeds ASVs’ sequences or OTUs’ representative sequences into a metric space where their pairwise distance can be utilized. The metric space embedding allows us to utilize their spatial structure to gain more detection power. To identify multi-scale signals, the MsRDB test adopts a propagation and separation (PS) approach (Polzehl and Spokoiny, 2000, 2006) to aggregate information from the neighborhood iteratively and adaptively. Notably, the MsRDB test only integrates OTUs/ASVs with similar levels of differential abundance so that we can capture the heterogeneous signal easily and avoid false discovery inflation caused by aggregation. As a generalization of robust differential abundance (RDB) test (Wang, 2023), the MsRDB test is also an iterative empirical Bayes method and thus inherits many good properties from the RDB test, including robustness to zero counts, compositional effect, and experimental bias. We evaluate the performance of the MsRDB test using both simulated and 16S rRNA data set from published studies. Through these numerical examples, we compare the MsRDB test with ASV-wise and taxon-wise analysis. We demonstrate that the MsRDB test can help reveal the multi-scale differentially abundant microbes at ASV resolution, providing more insights into the subtle difference between microbial communities.

## 2 Results

### 2.1 Overview of MsRDB Test

Before introducing the new MsRDB test, we first discuss why the results of differential abundance analysis highly rely on the choice of analysis unit. We focus primarily on the ASVs in this work for concreteness and illustration, while all the discussions and techniques introduced here readily apply to OTUs. The main idea in the taxon-wise analysis is to aggregate data of ASVs under the guidance of the assigned taxonomy. The way to aggregate ASV greatly impacts the results of differential abundance analysis. On the one hand, the aggregation usually enriches the differential abundant signal provided the ASVs assigned to the same taxon have similar levels of differential abundance ((i) and (iv) in Figure 2). On the other hand, when the ASVs have different levels of differential abundance, aggregation could lead to canceling out differential abundant signal ((ii) in Figure 2) and losing resolution ((iii) in Figure 2). However, the current taxonomy-based aggregation has no guarantee to ensure aggregation of ASVs with similar levels of differential abundance and may result in throwing out a considerable amount of unassigned ASVs. Can we design a new way of ASV aggregation that keeps all sequences in the analysis and only aggregate ASVs with similar levels of differential abundance?

**Figure 2:**
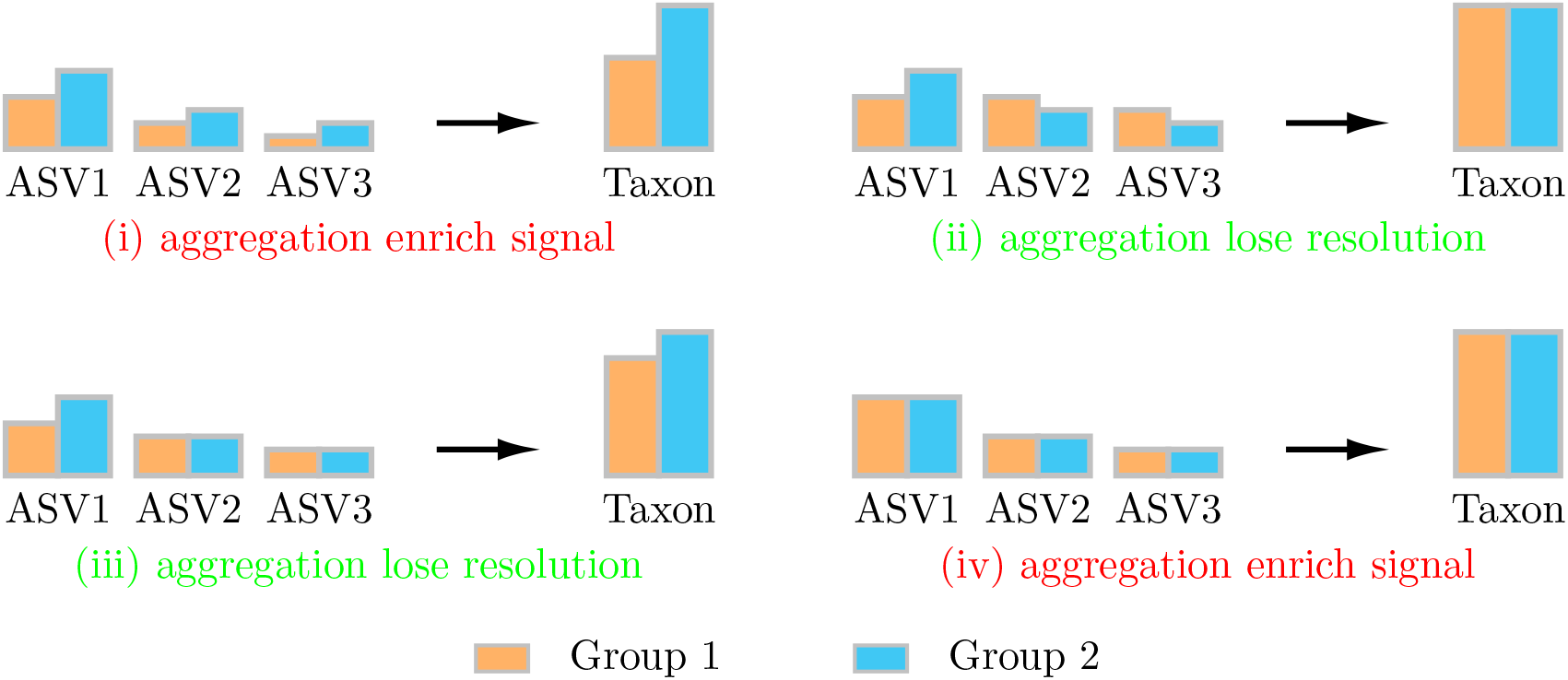
The effect in ASV aggregation. This figure illustrates four possibilities of ASV aggregation: (i) aggregate differential abundant ASVs with a similar level of differential abundance (e.g., all ASVs’ abundance in group 1 is smaller than abundance in group 2); (ii) aggregate differential abundant ASVs with different level of differential abundance (e.g., some ASVs’ abundance in group 1 are smaller than abundance in group 2 and some are larger); (iii) aggregate differential abundant ASVs and non-differential abundant ASVs; (iv) aggregate non-differential abundant ASVs. The aggregation in (i) and (iv) are beneficial as they can help enrich the signal, while the aggregation in (ii) and (iii) can make the analysis lose resolution and cancel out the signal. The bar plots in this figure represent absolute abundance.

We give a brief overview of the new MsRDB test, and a detailed description is explicated in the Supplementary Material. The MsRDB test consists of two main steps: 1) the ASV aggregation by a multi-scale and adaptive method (Figure 3C) and 2) differential abundance analysis on the weighted abundance table (Figure 3D). In the first step (Figure 3C), the MsRDB test embeds all ASVs’ sequences into a metric space so we can directly measure the similarities between any pair of ASVs’ sequences. Unlike the taxon-wise analysis, which measures ASVs’ similarity by assigned taxonomy, the idea of embedding into a metric space allows the MsRDB test to keep all sequences in the analysis. After embedding, the MsRDB test aggregates the data in each ASV’s neighborhood by assigning weights to each pair of ASVs *s* and *s*′

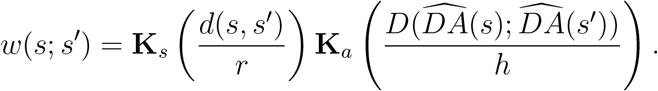

**Figure 3:**
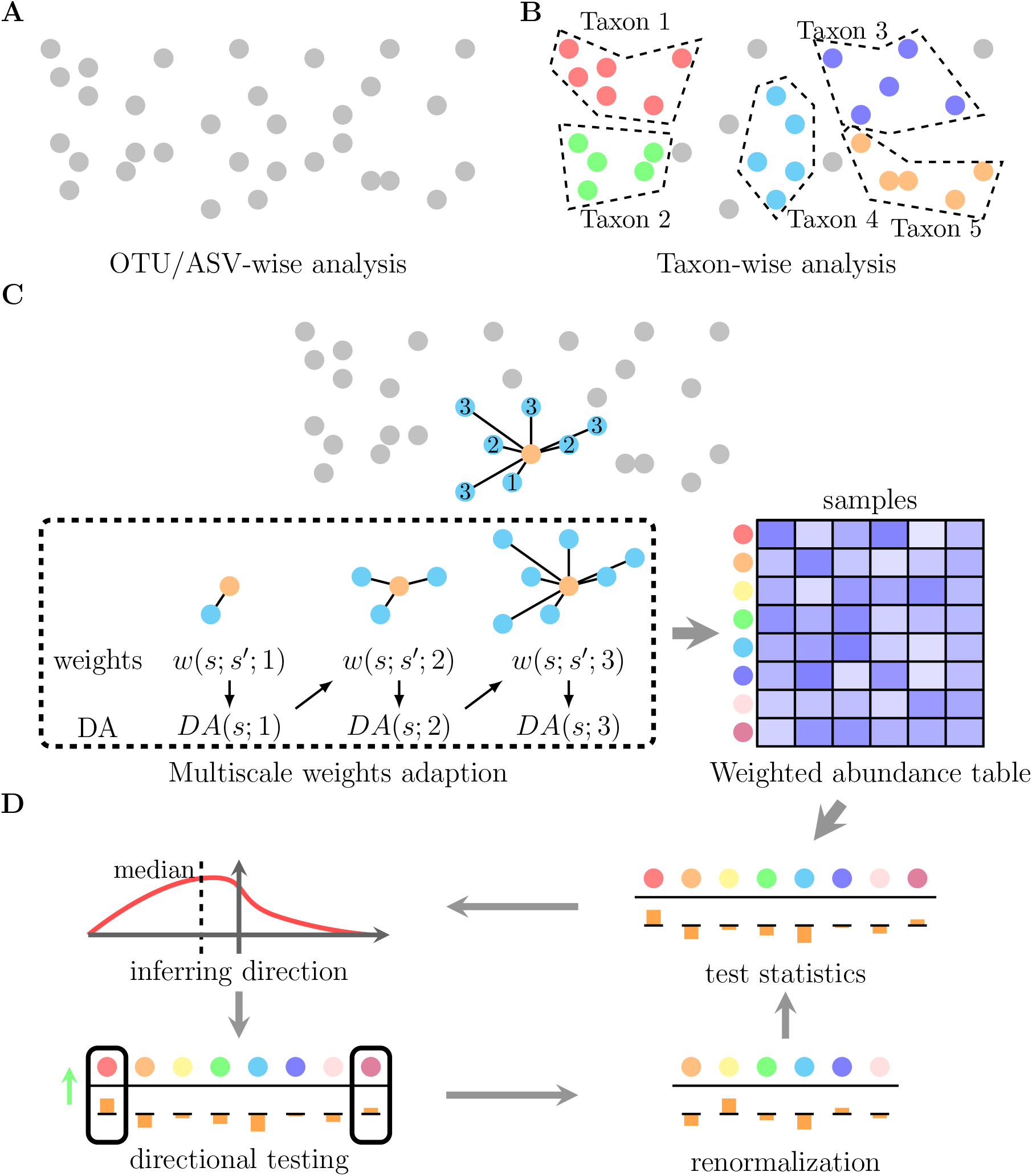
Illustration demonstrating OTU/ASV-wise analysis, taxon-wise analysis, and MsRDB test. (A) OTU/ASV-wise analysis treats OTUs or ASVs as independent units in differential abundance analysis. (B) According to the mapped or assigned taxon, OTUs or ASVs are first grouped as taxa. The taxa are regarded as independent units in taxon-wise analysis. (C) The first step in the MsRDB test constructs a weighted count table by exploiting the spatial structure of ASVs. Instead of independent units, the MsRDB test embeds the ASVs into a metric space and utilizes their spatial structure to gain more power. The PS approach used in the MsRDB test allows borrowing strength from the neighborhood in a multi-scale and adaptive way. (D) The second step in the MsRDB test identifies differential ASVs by iterative directional two-sample tests and renormalization.

The weights we use here reflect two types of information: how different the two ASVs’ sequences are and how different the levels of differential abundance are. Here, **K**_*s*_ is a kernel used to measure the sequence similarity, and **K**_*a*_ is designed to measure the similarity of differential abundance level. The differential abundance level in *w*(*s*; *s*′) is measured by a robust coefficient of estimated fold change

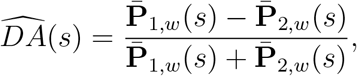

where 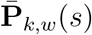 is the weighted mean of the relative abundance of *s* in the *k*th group, i.e., 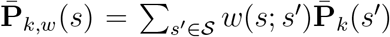, where 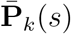 is the mean of the relative abundance of *s* in the *k*th group. Incorporating weights *w*(*s*; *s*′) in 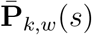 allows a more accurate estimation of differential abundance level but also raises a challenge in simultaneously evaluating weights and differential abundance level. Since *w*(*s*; *s*′) and 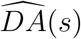 rely on each other, we adopt an iterative and multi-scale approach, called the propagation and separation method (PS), to update weights and the estimator for differential abundance level simultaneously and adaptively in a series of nested neighborhoods (Figure 3C). Through the PS approach, the weights in the MsRDB test allow each ASV to effectively borrow strength from other similar ASVs, thus increasing the power in detecting more subtle levels of differential abundance. Importantly, the weights in the MsRDB test reflect similarities of ASVs’ sequences and their differential abundance levels so that only ASVs with similar levels of differential abundance are aggregated. Based on the assigned weights, we aggregate the relative abundance of ASVs to obtain a weighted abundance table, where each row still corresponds to each ASV.

In the second step (Figure 3D), we conduct differential abundance analysis on the weighted abundance table obtained in the first step. Specifically, we formulate the differential abundance analysis in microbial compositional data as a reference-based hypothesis to account for the compositional nature of data and potential experimental bias (Brill et al., 2022). In the reference-based hypothesis, we do not directly test whether the relative abundance changes across groups, but we need to test if the fold change of each ASV is different from the majority of ASVs. To test such a hypothesis, we introduce a weighted version of the RDB test for the weighted abundance table, a generalized version of the vanilla RDB test (Wang, 2023). Like the vanilla RDB test, the weighted version RDB test also exploits the fact that the non-differential abundant ASVs share the same fold change, so the expectation of two-sample *t*-tests on them have the same signs. Using this fact, the MsRDB test adopts an iterative empirical Bayes framework to identify differential abundant ASVs. More concretely, the iterative empirical Bayes framework consists of two steps in each iteration: 1) integrates two-sample *t*-test statistics to infer the testing direction and then conduct directional two-sample *t*-test; 2) renormalizes the weighted abundance table with respect to the active ASVs (Figure 3D). Since only two-sample *t*-test statistics are evaluated, there is no need for an extra step to handle zero counts in the MsRDB test. Compared with the vanilla RDB test, the MsRDB test is designed for a weighted abundance table and uses ASVs’ taxonomy only for results interpretation instead of data aggregation.

In the MsRDB test, we need to specify two sets of tuning parameters: one is for the PS method, and the other is for the weighted RDB test. In the MsRDB test, we choose the tuning parameters of the PS method similar to the literature (Li et al., 2011; Wang et al., 2019). The performance of the MsRDB test is not very sensitive to most tuning parameters. The most sensitive parameter is the number of neighbors in the last iteration *k*. As shown in Section 2.2, the FDR may be slightly inflated when the number of neighbors *k* increases. In the weighted RDB test, we can choose the tuning parameters like the original RDB test. In particular, the most sensitive tuning parameter is the critical value of two-sample testing, which can be chosen by a BH-like procedure introduced in Wang (2023). See the detailed discussion in the Supplementary Material.

### 2.2 Borrowing Strength from Neighborhood can Increase Power

In this section, we study the numerical performance of the MsRDB test through a series of experiments on simulated data. These numerical experiments aim to characterize how borrowing the neighborhood’s strength can boost the power of the differential abundance test. In these simulation studies, we simulate the ASV data from a data set collected in Yatsunenko et al. (2012) (Supplement Material). With the simulated data set, we compare the false discovery rate (FDR) and power of the MsRDB test with different versions of the RDB tests and several state-of-art differential abundance tests.

The first set of simulation studies aims to investigate ASV aggregation’s effect by comparing different versions of RDB tests. In particular, we consider the vanilla RDB test, the *k*-nearest neighbors weighted RDB test, and the MsRDB test. The results of FDR and power are summarized in Figures 4, S1, and S2. These results suggest that the power is significantly improved if we incorporate information from the ASVs’ neighborhoods. In particular, the *k*-nearest neighbors weighted RDB test and the MsRDB test have larger power than the RDB test in all simulation experiments. When the number of neighbors increases, the power of both tests also increases since more information is borrowed from neighbors. However, simple incorporation from similar ASVs in the *k*-nearest neighbors weighted RDB test can lead to a highly inflated false discovery. Specifically, the FDR level can be as high as more than 50% sometimes. The inflation of false discovery is mainly because the non-differentially abundant ASVs can be falsely identified when they aggregate information from some differentially abundant neighbors. Unlike the RDB test and *k*-nearest neighbors weighted RDB test, the MsRDB test can help improve power while controlling FDR at the desired level since the PS approach only allows sharing information between ASVs with similar levels of differential abundance. Note that the FDR in the MsRDB test may be slightly inflated when the number of neighbors becomes larger. In conclusion, aggregating information from the neighborhood can increase power, but controlling false discovery needs smart aggregation.

**Figure 4:**
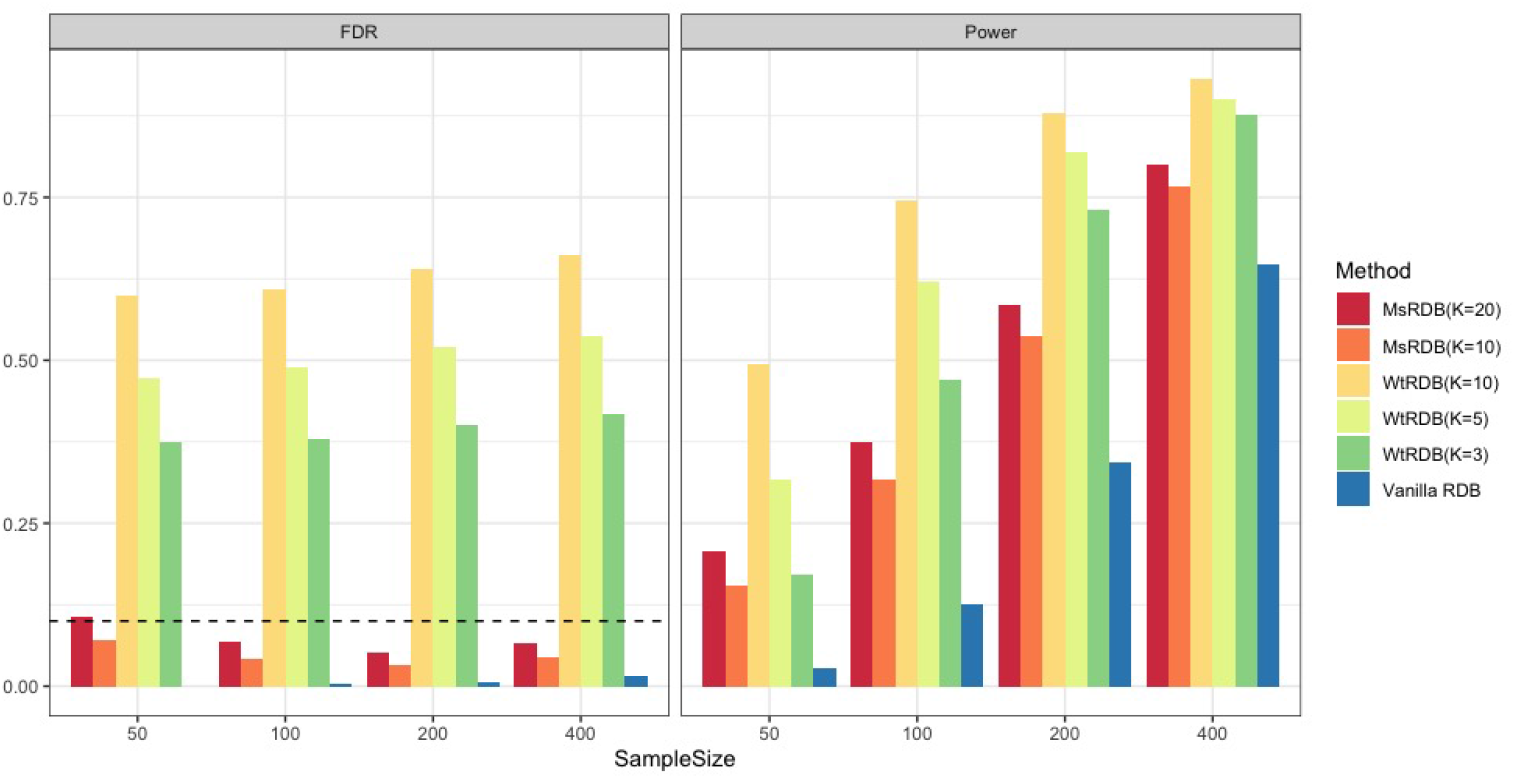
FDR and power comparisons of different versions of the RDB tests in simulated data sets. Left and right figures show the FDR and power of various versions of RDB methods when the sample sizes are 50, 100, 200, and 400. The dashed line in the left figure is the target level of FDR at 10%. The results in this figure show that aggregation of information from the neighborhood can increase the power. However, simple aggregation in k-nearest neighbors could result in inflation of false discoveries. In contrast, the MsRDB test increases the power and avoids FDR inflation since only ASVs with similar levels of differential abundance are aggregated.

These simulation experiments in Figures 4, S1, and S2 also help us investigate the influence of sample size, signal strength, and the number of differentially abundant ASVs on these three versions of the RDB tests. When the sample size increases, all these tests become more powerful and achieve similar FDR (Figure 4). Unlike the sample size, the number of differentially abundant ASVs has a relatively small influence on the performance of these tests (Figure S2). Similar to sample size, a large signal strength can lead to more powerful tests, but a small signal strength can slightly inflate the FDR in the MsRDB test (Figure S1). The intuition behind this phenomenon is that it is more difficult for the PS approach to distinguish ASVs with different levels of differential abundance when the signal strength is small. Hence, the non-differentially abundant ASVs are more likely to aggregate information from differentially abundant ASVs.

In the second set of simulation studies, we compare the MsRDB test with five state-of-art differential abundance tests, including the RDB test (Wang, 2023), ANCOM.BC (Lin and Peddada, 2020a), DACOMP (Brill et al., 2022), ALDEx2 (Fernandes et al., 2013), and StructFDR (Xiao et al., 2017). The performance of these six methods is evaluated by false discovery rate and power at the ASV level. All methods can control FDR at the ASV level very well, no matter the differential abundant ASVs belonging to different genera, classes, or local ASV clusters (Figure 5, S3, and S4). Due to ASVs aggregation, MsRDB and StructFDR are significantly more powerful than the other four methods in all experiments, consistent with the observation in the previous set of simulation studies. Compared with StructFDR, MsRDB has larger power as embedding ASVs into a metric space allows more flexibility in ASVs aggregation than using a phylogenetic tree. In addition, the MsRDB test can still control false discovery and maintain adequate power when the simulated ASV data has the measurement error and unbalanced sequencing depth (Figure S5 and S6).

**Figure 5:**
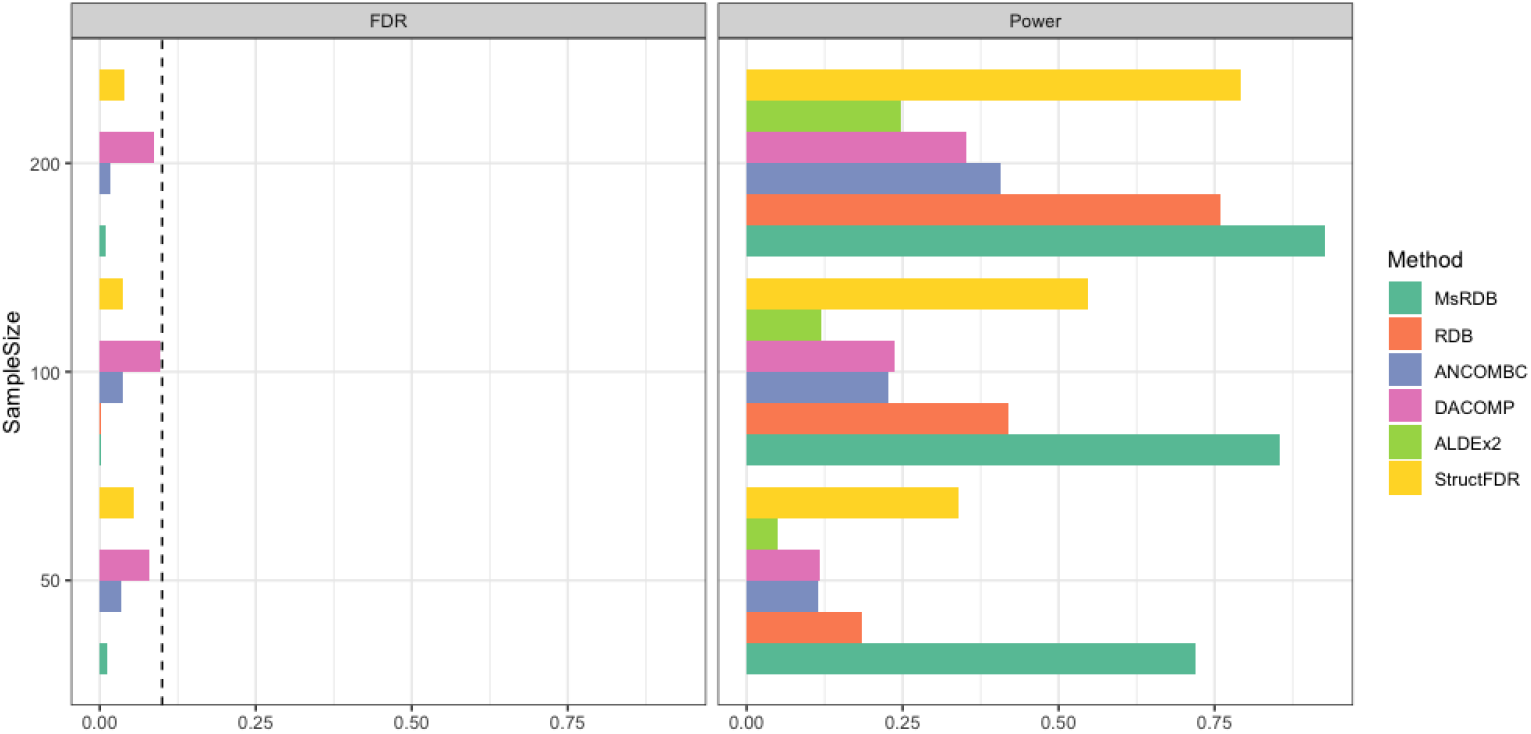
Comparisons of differential abundance tests in simulated data sets when differential abundant microbes are different genera. The differential abundant ASVs are all ASVs in genera *Ruminococcus, Bacteroides, Faecalibacterium, Oscillibacter*, and *Blautia*. Left and right figures show the FDR and power of differential abundance tests when the sample sizes are 50, 100, and 200. The dashed line in the left figure is the target level of FDR at 10%. All methods can control FDR at the ASV level very well. The results confirm that aggregating information from the neighborhood can lead to a more powerful test.

The last set of simulation studies aims to compare the performance of differential abundance tests at a resolution of chosen taxonomy rank, including genus and family. We consider six differential abundant tests: MsRDB, RDB with ASV as analysis unit, RDB with a genus (family) as analysis unit, ANCOM.BC with ASV as analysis unit, ANCOM.BC with a genus (family) as analysis unit, and StructFDR. A taxon is defined as a differential abundant taxon if at least one ASV belonging to this taxon is differential abundant. Unlike the previous experiments, the performance of the differential abundance test is evaluated by false discovery rate and power at the genus (family) level. Despite the analysis unit, all methods can control the FDR very well (Figure 6 and S7). We can observe that using the genus (family) as analysis units can lead to a more powerful test than using ASV when the sample size is small. It is also interesting to note that StructFDR and MsRDB can detect more differential abundant microbes than the other methods because of their ASV aggregating strategy and multi-scale technique. The three sets of simulation studies suggest that a proper aggregation strategy can improve the power of differential abundant tests.

**Figure 6:**
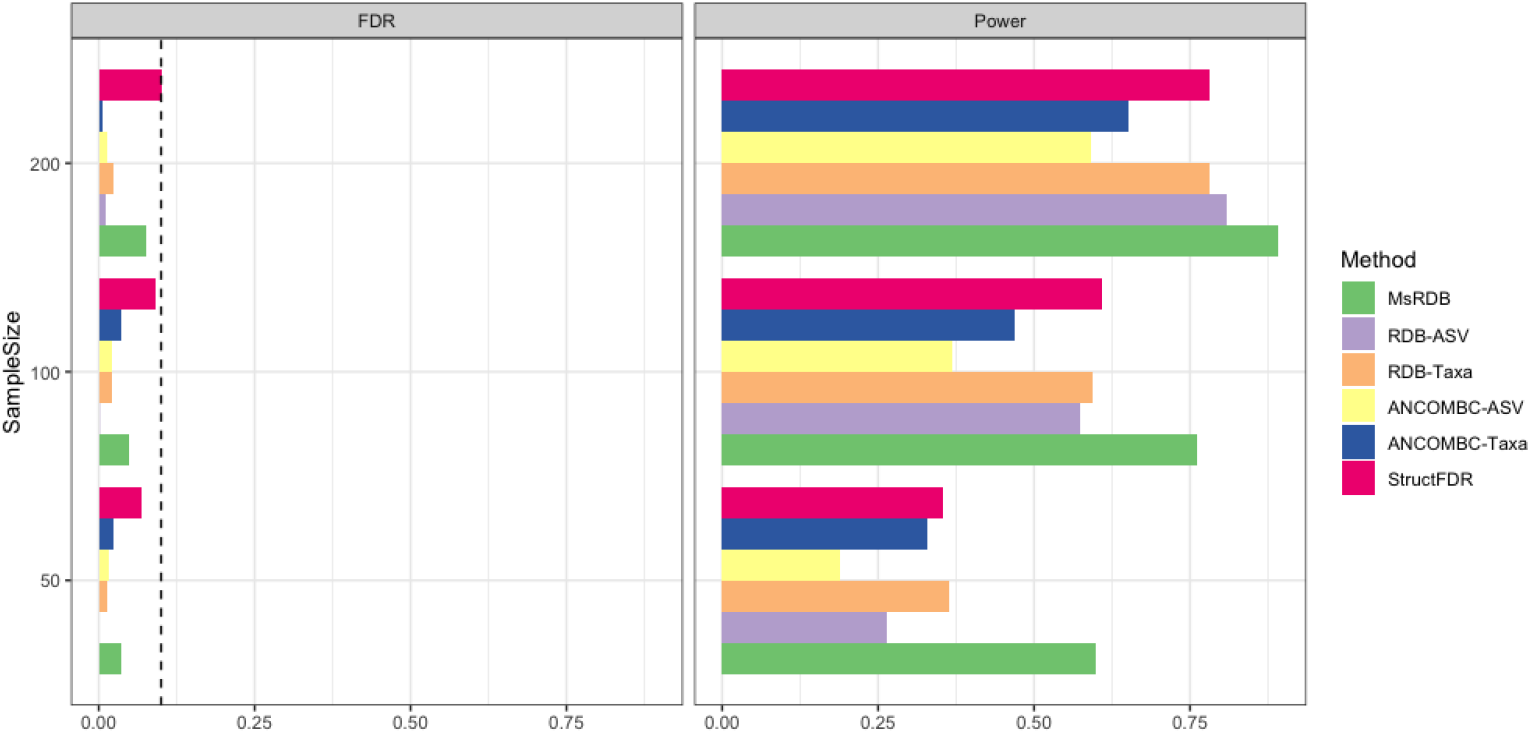
Comparisons of differential abundance tests in identifying differential abundant genera. The differential abundant genus is a genus with at least one differential abundant ASV. Left and right figures show the FDR and power of differential abundance tests when the sample sizes are 50, 100, and 200. The dashed line in the left figure is the target level of FDR at 10%. All methods can control FDR at the genus level very well. Choosing genus as analysis units can improve the power of RDB and ANCOM.BC when the sample size is small.

### 2.3 MsRDB Detects Differentially Abundant Microbes Associated with Immigration

To elucidate the differences between ASV-wise analysis, taxon-wise analysis at different ranks, and the MsRDB test, we apply these microbiome data analysis strategies to a gut microbiota data set from an immigration effect study (Vangay et al., 2018). In particular, we focus on the pairwise comparisons between Hmong female individuals who were living in Thailand (HmongThai, *n* = 96) or were born in the US but whose parents were born in Southeast Asia (Hmong2nd, *n* = 54), and European American female individuals (Control, *n* = 36). Through pairwise comparisons, we can identify the microbes associated with immigration. To conduct comparisons, we apply four different analysis strategies: ASV-wise analysis, taxon-wise analysis at the genus level, taxon-wise analysis at the class level, and the MsRDB test. In ASV-wise and taxon-wise analysis, we consider three differential abundance tests: ANCOM.BC (Lin and Peddada, 2020a), RDB test (Wang, 2023) and ALDEx2 (Fernandes et al., 2013).

Overall, the differential abundance analysis results suggest that Hmong2nd is very similar to Control while there are some differences between HmongThai and the other two groups. Regardless of the choices for differential abundance test, there are fewer discoveries if we regard each ASV as an analysis unit (Figure 7, S10, and S11). The low detection power in ASV-wise analysis is mainly because low counts at each ASV are not informative enough for an effective comparison. The power is significantly improved after aggregation based on taxonomy, but we still need to choose an appropriate rank for analysis. A lower taxonomy rank brings finer resolution but also results in throwing out a fair amount of sequence. Specifically, at least 5% of valuable sequences in more than half samples are assigned to the NA genus and thus removed from the analysis if we choose genus as analysis unit (Figure 7D). Compared with ASV-wise analysis, the genus-wise analysis leads to more discoveries. On the other hand, if a higher taxonomy rank is used, we can keep more sequences but could lose resolution. More concretely, only a tiny proportion (< 1% in most samples) of sequences are removed from the class-wise analysis (Figure 7D). However, the class-wise analysis only reports differentially abundant microbes at the class level and loses the resolution. In particular, we can observe that the analysis at different ranks contradicts each other sometimes in Figure 7. For example, genus-wise analysis suggests that some genera in class *Coriobacteriia, Bacilli, Desulfovibrionia, Negativicutes*, and *Gammaproteobacteria* are differentially abundant in HmongThai and Control comparison, but class-wise analysis makes an opposite conclusion.

**Figure 7:**
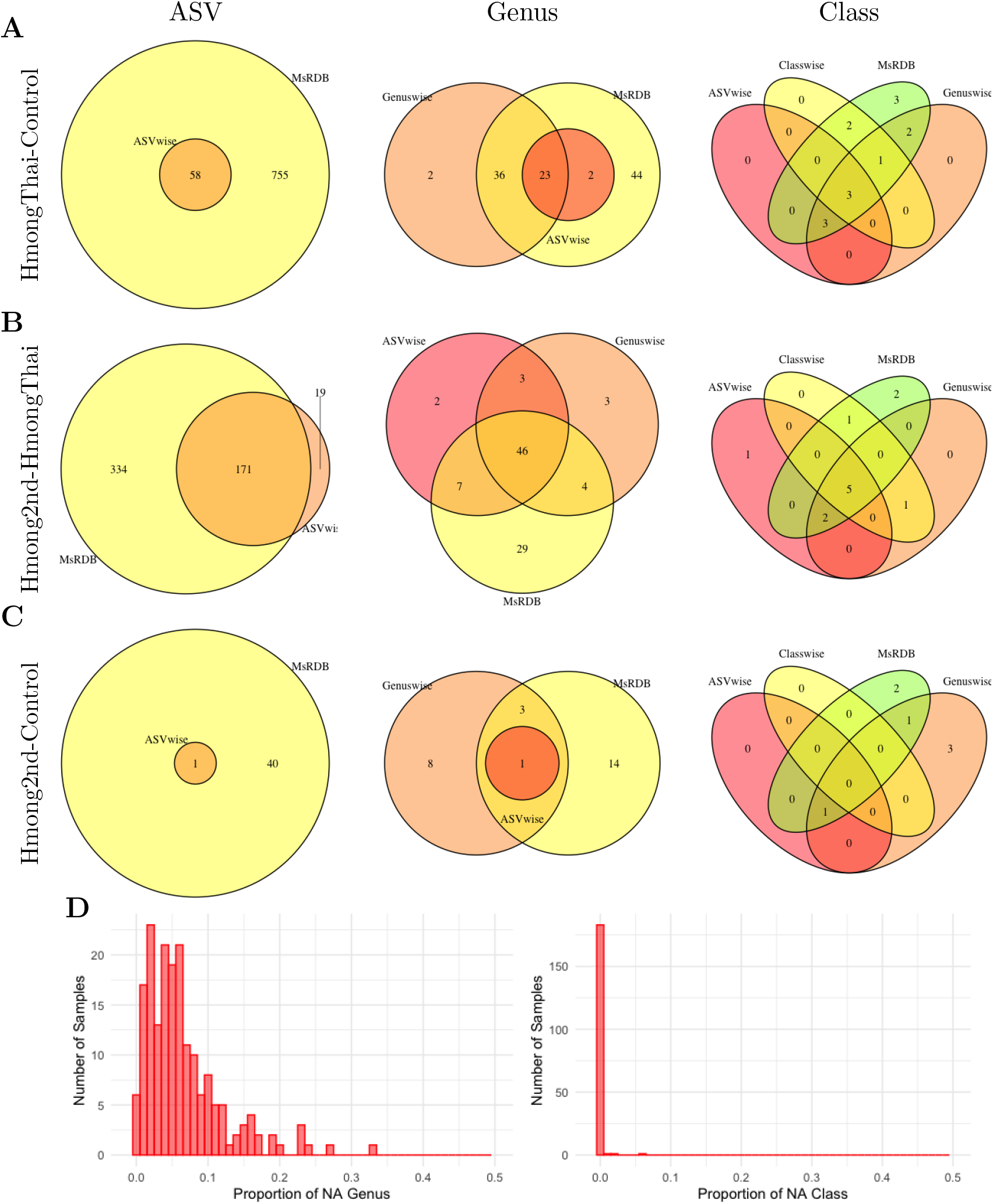
Comparisons between ASV-wise analysis, taxon-wise analysis, and MsRDB test on the microbiota data set in an immigration study. (A-C) shows the overlap of results obtained by different analysis strategies in pairwise comparisons of three groups of people. The three rows are the comparisons of HmongThai vs. Control, Hmong2nd vs. HmongThai, and Hmong2nd vs. Control. The first column is the overlap of identified ASV by ASV-wise analysis and the MsRDB test. The second column is the overlap of identified genera by ASV-wise analysis, genus-wise analysis, and the MsRDB test. The third column is the overlap of identified classes by ASV-wise analysis, genus-wise analysis, class-wise analysis, and the MsRDB test. (D) The histograms of proportions of NA genus (left) and class (right) in each sample. Most samples need to remove at least 5% sequences from the genus-wise analysis.

The MsRDB test combines the advantages of both ASV-wise analysis and taxon-wise analysis. Like ASV-analysis, the MsRDB test utilizes all sequences and detects the differentially abundant microbes at ASV resolution. Since strength is borrowed from the neighborhood, the MsRDB test is more powerful and has more discoveries than ASV-wise or taxon-wise analysis. Importantly, most discoveries by all different methods are recovered by the MsRDB test, whether they are detected at ASV, genus, or class level (Figure 7, S10, and S11). This phenomenon is mainly because the weights in the MsRDB test are constructed iteratively so that the MsRDB test can capture multi-scale signals.

The aggregation strategy in the taxon-wise analysis can help increase power but could also miss some heterogeneous signals. To illustrate this, we focus on the comparison between group HmongThai and Control. The class-wise analysis suggests that *Bacilli* is a non-differentially abundant class (Figure 8A). However, *Bacilli* is known to be associated with the western diet and is expected to be more abundant in the Control group (Clarke et al., 2012). Through the genus-wise analysis, we identify five differentially abundant genera in *Bacilli* (Figure 8C). Figure 8A shows the box plot of the abundance in these five significant genera and the rest of the genera in *Bacilli*. These dominant non-differentially abundant genera make it difficult to identify *Bacilli* in class-wise analysis. This example indicates we might miss some subtle heterogeneous signals in taxon-wise analysis.

**Figure 8:**
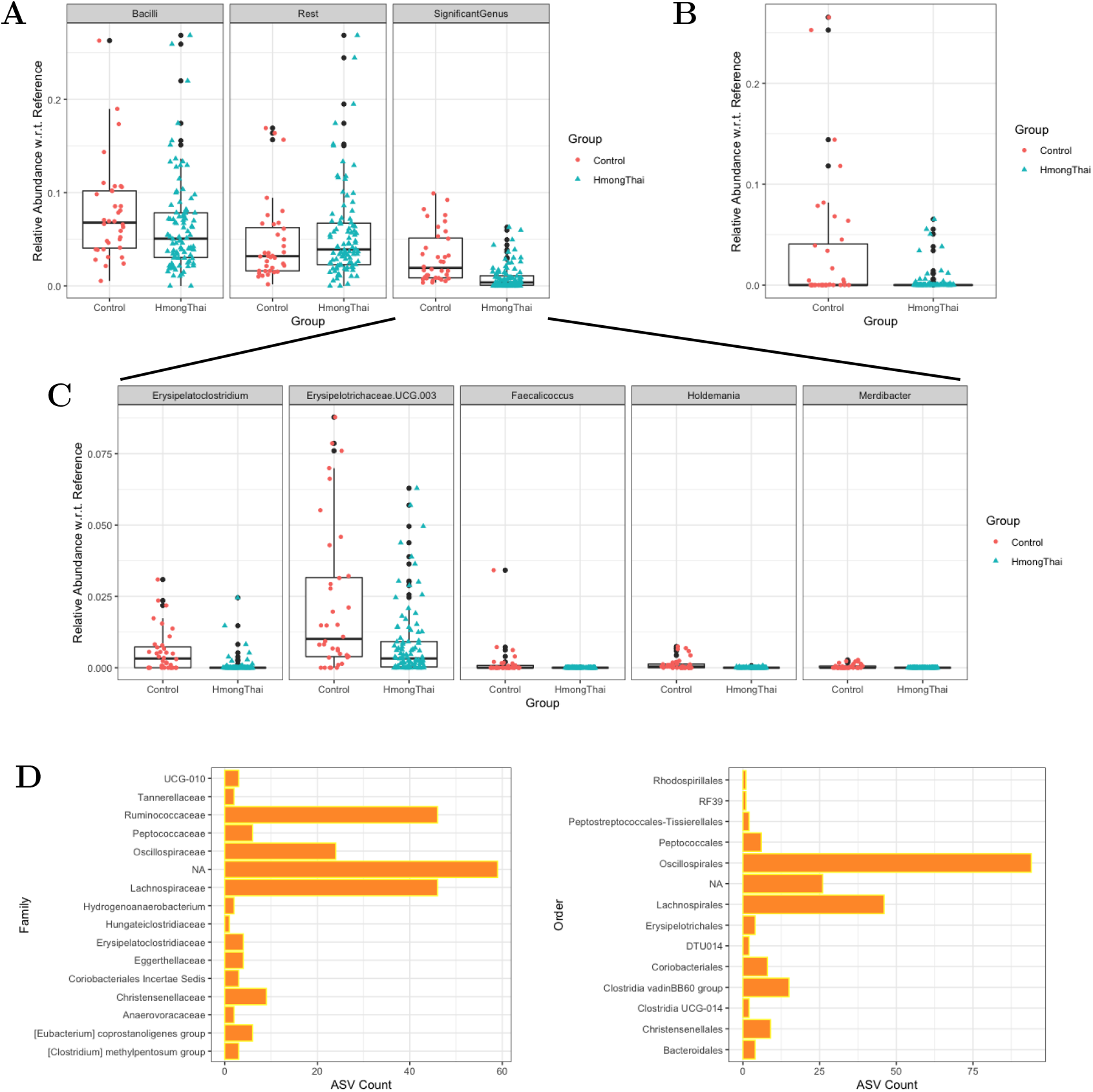
Taxa detected by different differential abundance analysis strategies on the real microbiota data set. The reference set in all box plots is the collection of all non-differentially abundant ASVs revealed by the MsRDB test. (A) The box plots show the relative abundance with respect to the reference set in class *Bacilli* (left), in five significant genera within *Bacilli* (right), and in the rest of genera within *Bacilli* (middle). (B) The box plot shows the relative abundance with respect to the reference set in significant ASVs within genus *Phascolarctobacterium*. (C) The relative abundance with respect to the reference set in five significant genera within *Bacilli* is shown. The five significant genera are identified by genus-wise analysis. (D) The family (left) and order (right) of significant ASVs assigned to the NA genus. These ASVs are identified by the MsRDB test and removed from the genus-wise analysis.

Unlike taxon-wise analysis, the MsRDB test only aggregates ASVs with similar levels of differential abundance and thus can easily capture these heterogeneous signals. This special aggregation strategy partially explains why the MsRDB test identifies more genera or classes than taxon-wise analysis and potentially provides more biological insights into the microbial communities. In MsRDB analysis results, several ASVs assigned to genus *Phascolarctobacterium* are more abundant in group Control than HmongThai (Figure 8B, present in 26% of HmongThai < present in 47% of Control), while it is non-differentially abundant in genus-wise analysis. This discovery is consistent with previous results that *Phascolarctobacterium* is associated with a high-fat diet (Ariefdjohan et al., 2017). MsRDB analysis results suggest a decrease in the abundance of several ASVs assigned to *Ruminococcus torques* in HmongThai, compared with Control (Figure S9A, present in 21% of HmongThai < present in 78% of Control). A plant-based diet in Southeast Asia could explain the decrease in the abundance of *Ruminococcus torques* (Meslier et al., 2020). It is also interesting to observe that the rest of the ASVs assigned to *Ruminococcus torques* are not significantly differentially abundant, so genus-wise analysis finds it difficult to detect.

The ASVs removed from the taxon-wise analysis could also help to provide insight into the microbial communities. The MsRDB test identifies a few significant ASVs removed from genus-wise analysis in HmongThai and Control comparison. Figure S9B compares these significant ASVs’ abundance if we aggregate them together. There is a significant increase in Control compared with HmongThai. Although these significant ASVs are assigned to the NA genus, their taxonomy classification at family or order rank can still provide a rich source of information (Figure 8D). Specifically, these genus-unassigned ASVs mainly come from the family *Ruminococcaceae, Oscillospiraceae*, and *Lachnospiraceae*, or order *Oscillospirales* and *Lachnospirales*.

### 2.4 MsRDB Reveals Microbial Biogeography of Wine Grapes

To further demonstrate the performance of the MsRDB test, we apply it to a grape microbiota data set (Bokulich et al., 2014). We consider studying the association between growing region and microbial community of grape Chardonnay. The grape must samples we consider here are divided into three groups based on their growing regions: 19 samples from Sonoma, 46 samples from Napa, and 11 samples from Central Coast. Unlike the previous data set, a considerable amount of sequences in this data set are assigned to the NA genus and family (Figure 9A). We need to remove these sequences from analysis if we adopt taxon-wise analysis to detect differentially abundant microbes at a resolution of genus or family. We apply the MsRDB test and ASV-wise analysis to explore differentially abundant microbes at an ASV resolution. Again, the MsRDB test is more powerful and can help detect more significant ASVs (Figure S12).

**Figure 9:**
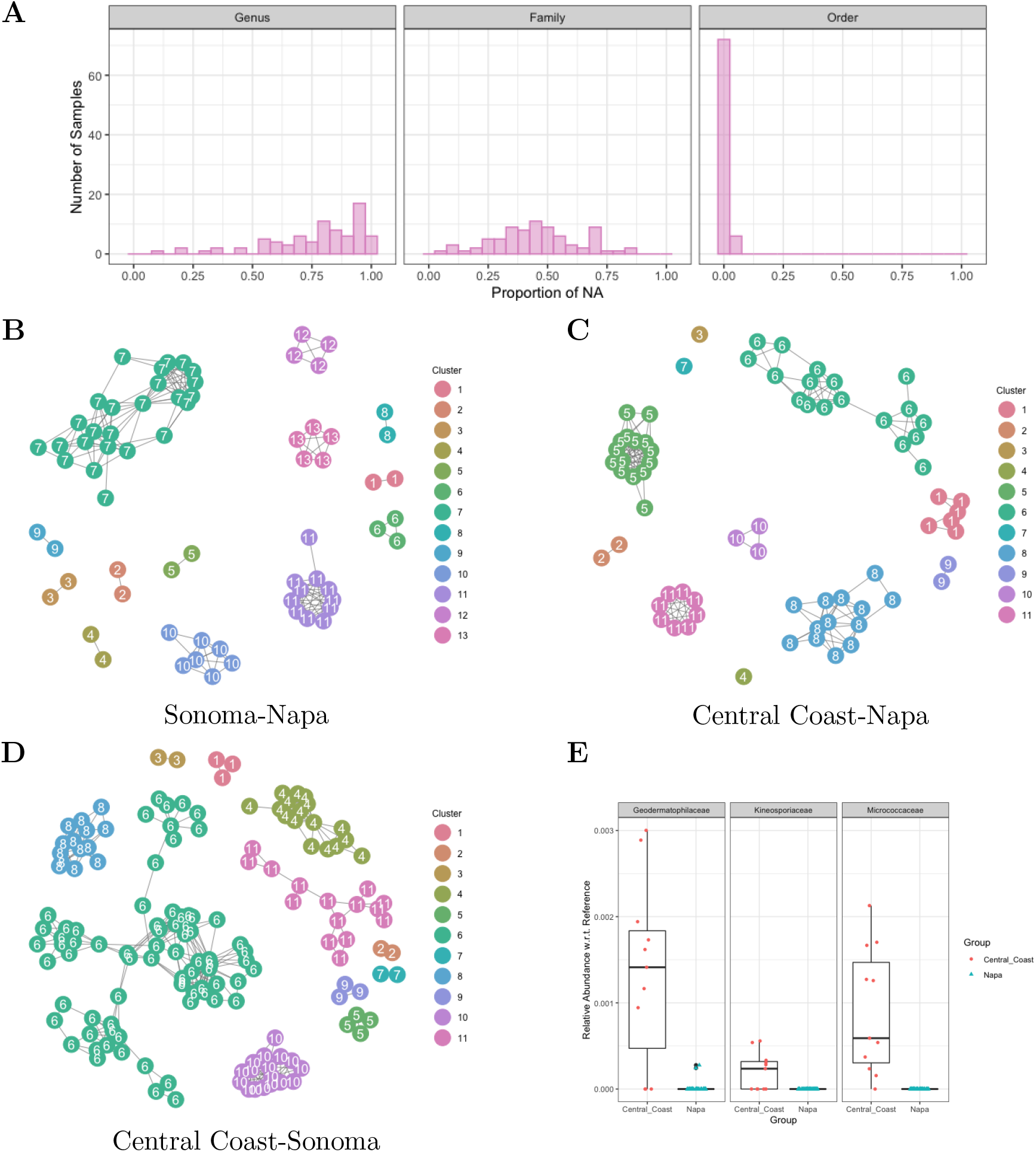
Analysis of the grape microbiota data set by MsRDB test. (A) The histograms of proportions of the NA genus (left), class (middle), and order (right) in the grape microbiota data. A considerable amount of sequences are removed from the genus-wise or family-wise analysis. (B-D) The clustering results of significant ASVs in the pairwise comparisons. 13 clusters are obtained in Sonoma vs. Napa comparison, 11 clusters are obtained in Central Coast vs. Napa comparison, and 11 clusters are obtained in Central Coast vs. Sonoma comparison. (E) The box plots show the relative abundance with respect to the reference set in clusters 8, 9, and 10 in Central Coast vs. Napa comparison.

We cluster the differentially abundant ASVs into small groups since only partial information on the microbial taxonomy is known in this data set. Figures 9B-D show the clustering results in pairwise comparisons between Sonoma, Napa, and Central Coast and indicate that the differentially abundant microbes naturally form around 10 small groups in each comparison. After comparing each small group with the assigned taxonomy, the small groups can correspond to different taxonomy ranks (Table S1). For example, in comparing between Central Coast and Sonoma, ASVs in cluster 1 come from the same genus *Lactobacillus* while ASVs in cluster 11 come from the same order *Burkholderiales*. This example again demonstrates that the MsRDB test can indeed capture multi-scale signals. Although some ASVs lack assigned genus and family, they may still provide insight into microbial communities. For instance, ASVs cluster 8 in the Sonoma and Napa comparison are assigned to NA genus and family, but the information in order *Enterobacterales* is still beneficial. The analysis in (Bokulich et al., 2014) concludes that class *Actinobacteria* is more abundant in Central Coast than Napa. Our results confirm this finding and further suggest that the differentially abundant microbes come from family *Micrococcaceae* (cluster 8), *Geodermatophilaceae* (cluster 9), and *Kineosporiaceae* (cluster 10) (Figure 9E).

## 3 Discussion

Differential abundance analysis is a challenging and complicated statistical problem because the observed data set in microbiome studies only reflects the relative abundance of taxa, has excessive zeros, and is perturbed in experiments. Besides these major challenges, this work focuses on another important practical issue in differential abundance analysis: the choice of analysis unit. Through several numerical examples, we show that the analysis unit has a significant impact on the results of differential abundance analysis. On the one hand, the ASV-wise analysis can help detect differentially abundant microbes at the finest resolution but usually has small detection power. On the other hand, taxon-wise analysis enforce a tradeoff between resolution and sequence utility, although it can help increase detection power.

MsRDB test has both advantages of ASV-wise and taxon-wise analysis: it detects the differentially abundant microbes at a resolution of ASV, has larger detection power, and keeps all sequences in the analysis. Due to these good properties, the MsRDB test not only recovers most discoveries detected by conventional ASV-wise or taxon-wise analysis but also identifies some extra subtle differences between microbial communities. These new findings revealed by the MsRDB test could potentially provide more biological insights into the microbial communities and help identify new diagnostic or prognostic biomarkers in disease. As a generalization of the RDB test, findings reported by the MsRDB test is reliable and trustworthy as it is robust to compositional constraint, prevalent zero counts and experimental bias in the microbial data set. We hope this new differential abundance analysis method will help biological researchers in making more scientific discoveries.

This work not only develops a new differential abundance test but also suggests a change in how we utilize the ASVs’ sequence information in the differential abundance analysis. Conventional differential abundance analysis treats ASVs (or taxa) as independent units and thus does not take the most advantage of sequences’ information. In contrast, we show that embedding into a metric space makes it possible to exploit the spatial structure among ASVs’ sequences and significantly improve the power of differential abundance analysis. The MsRDB test presented in this work is mainly designed based on the RDB test because of its robustness. However, the perspective of embedding and spatial analysis is readily generalizable to other popular differential abundance analysis methods. For instance, after embedding into a metric space, the linear models used in ANCOM.BC can also work with the PS approach, as illustrated in Li et al. (2011). Therefore, we believe this new perspective will lead to more exciting methods for analyzing microbial compositional data.

The current MsRDB test mainly considers a standard two-sample *t*-test as test statistics and Gaussian distribution as its null distribution. As we illustrate in our numerical examples, such a strategy works very well when the sample size is large. However, when the sample size becomes small, the false discovery might not be controlled well. This problem might be alleviated by a rank-based test, such as the Wilcoxon test, as the null distribution of the rank-based test does not rely on the underlying distribution. Because of the two-sample *t*-test, the current MsRDB test is designed for testing the mean shift between groups. It would also be interesting to explore if the framework of the MsRDB test can be used to test other characteristics of microbial distribution, such as dispersion. Another potential extension of the MsRDB test is to study the association between microbial community and a continuous or multiple-level outcome, e.g., BMI. This extension is possible if we consider replacing the *t*-test with a correlation test, as Wang (2023) illustrates.

## 4 Methods

### 4.1 Model for Microbiome Data

In this work, we consider the ASV counts data from *m*_1_ + *m*_2_ subjects, where *m*_1_ subjects are in the first group, and *m*_2_ are in the second group. Let 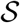 be the collection of observed distinct ASVs, and 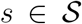 represents an ASV. For the *j*th subject in the *k*th group, we observe a vector of ASVs’ counts 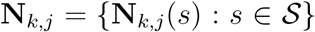, where **N**_*k,j*_(*s*) is the number of ASV *s* observed in the subject, and a vector of covariates **X**_*k,j*_, such as age and gender. Since the total counts of ASVs observed from each subject do not reflect the absolute abundance, we usually normalize the count data from each subject as a vector of observed relative abundance 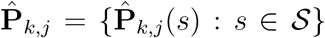, where 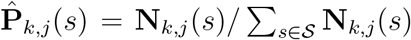. In practice, such a vector of observed relative abundance is also called a compositional data vector. In differential abundance analysis, our goal is to compare the observed relative abundance from these two groups of subjects.

We assume the ASV counts data are drawn from the following model. For the *j*th subject in the *k*th group, there exists a latent vector of absolute abundance 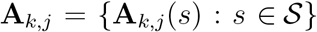, which we do not observe in practice. We assume the absolute abundance vectors and covariates vectors are drawn from two populations

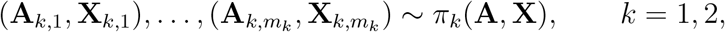

where **X** ∈ ℝ^*d*^. *d* = 0 means that there are no observed covariates in the data set. These absolute abundance vectors **A**_*k,j*_ represent the abundance of ASVs in each subject. Our observed ASV counts data **N**_*k,j*_ are related to absolute abundance **A**_*k,j*_ through a relative abundance model with experimental bias. Specifically, we adopt the multiplicative factors model (McLaren et al., 2019) for experimental bias in relative abundance

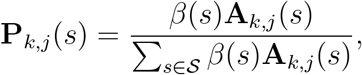

where *β*(*s*) is the measurement efficiency factor for the ASV *s*, which could related to variation in multiple technical factors, such as rRNA extraction efficiency(Olson and Morrow, 2012) and PCR primer binding preference (Brooks et al., 2015). *β*(*s*) is used to characterize the taxa specific bias and usually unknown in advance. Here, we want to emphasize that the relative abundance **P**_*k,j*_(*s*) already takes the experimental bias into account. Given the relative abundance, we assume the ASV counts **N**_*k,j*_ is drawn from a multinomial model

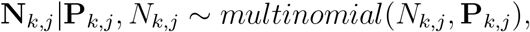

where *N_k,j_* is the total reads we observe for each subject. This model suggests that the observed relative abundance 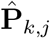 is just an empirical version of relative abundance **P**_*k,j*_.

### 4.2 Reference-Based Hypothesis

We need to define a rigorous statistical hypothesis in differential abundance analysis. Although it seems that the definition of a hypothesis is straightforward, compositional constraints and experimental bias make it challenging to define a statistically identifiable and interpretable hypothesis. See more discussions in Wang (2023). The literature usually defines the statistical hypothesis based on a parametric model, but it might not be easy to interpret when the model is misspecified. This work defines a rigorous statistical hypothesis without assuming a parametric model. Specifically, we formulate the differential abundance analysis as a reference-based hypothesis testing problem (Brill et al., 2022).

We first introduce the concept of the reference set (Sohn et al., 2015; Brill et al., 2022). A subset of ASVs, 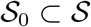, is called a reference set if 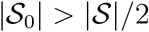 and there exists a constant *b* such that

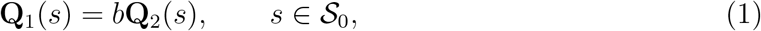

where 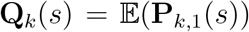 is the expected relative abundance of ASV *s* in the *k*th group. Here |·| is the number of elements in the set. The definition of the reference set suggests that the expected relative abundance of ASVs in the reference set changes in the same amplitude across groups. In other words, the fold changes of ASVs’ relative abundance in the reference set are the same. We define the reference set as (1) because the relative abundance changes caused by compositional effect and experimental bias alone have the same amplitude. For example, in Figure 1, the relative abundance of orange, green, and blue components change across groups due to the compositional effect and experimental bias alone, but their relative abundances change in the same amplitude (37.5%/30%=18.75%/15%=6.25%/5%=1.25).

After introducing the reference set, we now define the reference-based hypothesis. The idea of the reference-based hypothesis is to consider the reference set as a benchmark and compare other ASVs with this benchmark. Specifically, we consider the following reference-based hypothesis at ASV *s*

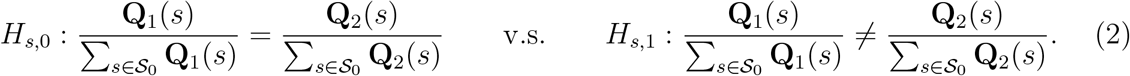

Since (2) is defined for each ASV, testing the above hypothesis can identify the differential abundant microbes at ASV resolution. We can also write the hypothesis in (2) as the following equivalent form

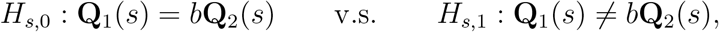

where *b* is defined in the reference set. This above definition suggests that the reference-based hypothesis does not directly test whether the relative abundance changes across the two groups or not but aims to test if the fold change of each ASV is different from the majority of ASVs due to the assumption 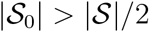. In the reference-based hypothesis, a change in the differential components can be interpreted as a change with respect to the collection of all non-differentiate ASVs. How does this reference-based hypothesis relate to the hypothesis defined by absolute abundance? If there exists a set of non-differential abundant ASVs (defined by the absolute abundance), this set of ASVs can be seen as a reference set. So testing the above reference-based hypothesis is roughly equivalent to directly testing if there is a change in absolute abundance.

A toy example is shown in Figure 1 to illustrate the idea of the reference set and reference-based hypothesis. In Figure 1, we have two ecosystems of red, orange, green, and blue components, *A* and *B*, and their absolute abundances are shown on the left-hand side. Due to the sampling process and experimental bias, the observed counts and relative abundance, shown on the right-hand side, cannot honestly reflect the relative relationship between different components’ absolute abundance. However, despite the experimental bias, the reference-based hypothesis framework suggests that the orange, green, and blue components can be seen as a reference set since their relative abundances change in the same amplitude (37.5%/30%=18.75%/15%=6.25%/5%=1.25). Based on this reference set, only the red component belongs to the alternative hypothesis as its fold change differs from the other three (37.5%/50% ≠ 1.25), and the orange, green, and blue components belong to the null hypothesis. These conclusions are consistent with the fact that there is only a change in the red component’s absolute abundance. This toy example shows that the reference-based hypothesis is a statistically identifiable and interpretable hypothesis for differential abundance analysis.

### 4.3 Multiscale Adaptive Robust Differential Abundance Analysis (MsRDB)

To test the reference-based hypothesis in (2), we introduce a multi-scale adaptive robust differential abundance test (MsRDB). In the MsRDB test, the input is the ASV counts table and the corresponding ASVs’ sequences. Given the input, the roadmap of the MsRDB test consists of the following three parts:

1. Initialization: the ASV counts table is transformed into the relative abundance table. The ASVs’ sequences are used to evaluate the ASV distance matrix. The details of the ASV distance matrix are explained in Section 4.3.1.
2. Weights calculation: evaluate weights *w*(*s*; *s*′) by applying propagation and separation approach to ASV relative abundance table and ASV distance matrix. The output of this step is a weighted ASV table. The details are explicated in Section 4.3.2.
3. Differential abundant ASV detection: identify differential abundant ASVs by applying the weighted RDB test to the weighted ASV relative abundance table. The details are explicated in Section 4.3.3.

Through these three parts, the MsRDB test identifies a collection of differentially abundant ASVs. Note that the analysis of the MsRDB test does not need information taxonomy, while the interpretation results may still need it. For the choices of tuning parameters in this algorithm, see Section 4.4.3.

#### 4.3.1 Embedding of ASVs’ Sequences

Different ASVs are not equally distinct, as some are more similar than others. Motivated by this observation, we can consider a general way to incorporate the similarity between ASVs. Specifically, we opt to embed different ASVs into a metric space and define a distance between ASVs’ raw sequences *d*(·,·). To reflect different aspects of ASVs, we provide several different possible ways to calculate the distance between two distinct ASV sequences:

- *Pairwise alignment distance*. One commonly used way to compare two sequences is pairwise sequence alignment, e.g., the Needleman-Wunsch algorithm. After alignment, the scoring schemes in the alignment algorithm can be used to calculate the distance between sequences.
- *kmer-based distance*. Another way to compare two sequences is *k*mer-based distance. Given a positive integer *k*, each sequence is transformed to a vector of *k*mer counts. Then, a distance between two *k*mer count vectors is used to measure the dissimilarity of sequences. For example, Euclidian squared, and Mahalanobis distances are two of the most commonly used distances (Gentleman and Mullin, 1989).
- *Phylogenetic distance*. If the evolutionary history of sequences is believed to relate to the outcome of interest, we can also use the existing phylogenetic tree to measure the dissimilarity of sequences. Specifically, the sequences are placed on a reference phylogenetic tree by phylogenetic placement algorithms, such as SEPP (Janssen et al., 2018). The distance between the sequences can then be defined as the length of the unique path connecting them on the tree.

This list of distance options could be incomplete, as some other information can also be incorporated into the distance. For example, the sequence can also be used to predict gene families or pathways (Douglas et al., 2020), and they can define the distance between sequences. In addition, a weighted sum of several distances could also be used when needed. In most of the analyses in this work, the distances between ASVs are calculated by function `DistanceMatrix` in `DECIPHER` R package (Wright, 2016). The output of this step is a distance matrix 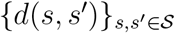.

#### 4.3.2 Multiscale Adaptive Weights Construction

In taxon-wise analysis, ASVs are aggregated by assigned taxonomy since the ASVs assigned to the same taxon are believed to be similar. Instead of using taxonomy, we assign weights *w*(*s, s′*) to each pair of ASVs to capture their similarity. Compared with taxonomy-based aggregation, the weights provide more flexibility in ASV aggregation and avoid throwing out the unassigned sequence in taxonomic classification. The general goal of weights construction is to increase detection power by borrowing strength from the neighborhood and reduce false discoveries by avoiding aggregation of differential and non-differential abundant ASVs.

##### Form of weights

One important advantage of weights *w*(*s, s′*) is that they can incorporate different types of similarity information between ASVs. More specifically, the weights we use here reflect two types of information: how different the two ASVs’ sequences are and how different the levels of differential abundance are. Compared with the classical form of weights in nonparametric statistics, our weights also reflect the similarity of differential abundance levels. The rationale for incorporating differential abundance levels is to ensure ASVs aggregation with similar levels of differential abundance. Specifically, the form of weights we consider is

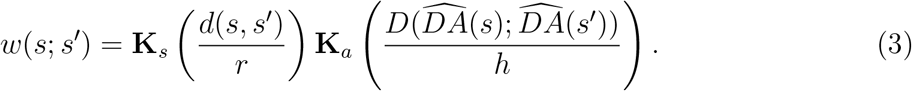

Here, **K**_*s*_ and **K**_*a*_ are two non-negative and non-increasing kernels with compact support, e.g., **K**_*a*_(*x*) = max(1 – *x*, 0) and **K**_*s*_(*x*) = **I**(*x* < 1), where **I** is an indicator function. **K**_*s*_ and **K**_*a*_ are used to incorporate different types of information:

- **K**_*s*_ is a kernel used to measure sequence similarity. In the kernel **K**_*s*_, *d*(*s,s′*) is the distance between ASVs’ sequences introduced in Section 4.3.1, and *r* is a tuning parameter to control the size of the neighborhood. We only aggregate the information from the ASVs which share similar sequences through the kernel **K**_*s*_.
- **K**_*a*_ is a kernel used to measure differential abundance level similarity. The reference-based hypothesis suggests that the level of differential abundance can be captured by fold change across the groups. We introduce a robust coefficient of fold change to measure the level of differential abundance

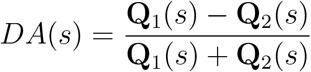

where 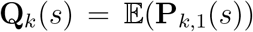 is the expected relative abundance of ASV *s* in the *k*th group. All non-differential abundant ASVs have *DA*(*s*) = (*b* – 1)/(*b* +1) and thus share the same level of differential abundance, where *b* is the constant in the definition of the reference set. Since *DA*(*s*) cannot be measured directly, we introduce the corresponding estimator of *DA*(*s*)

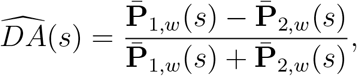

where 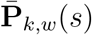 is the weighted mean of the relative abundance of ASV *s* in the *k*th group, i.e., 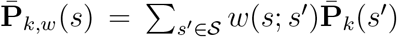. Here 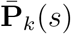 is the mean of the relative abundance of ASV *s* in the *k*th group. To compare the differential abundance level, we define the following distance between *s* and *s*′

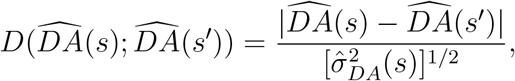

where 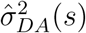 is an estimator for the variance of 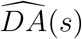. *h* in **K**_*a*_ is a tuning parameter to penalize the similarity of differential abundance between ASVs.

##### Propagation and separation

There are two challenges to evaluating the weights in (3): 1) the definition of *w*(*s*; *s*′) and 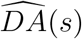 rely on each other, so we need to evaluate them simultaneously; 2) the performance of the weighted method usually depends on the choice of neighborhood size *r* and the optimal choice of *r* could be different at different ASVs. To alleviate these challenges, we adopt an iterative and multiscale approach, propagation and separation method (PS) (Polzehl and Spokoiny, 2000, 2006), to select neighborhood size and update estimation for *w*(*s*; *s*′) and 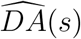 simultaneously. The PS approach has been widely used in various imaging analyses (Li et al., 2011; Wang et al., 2019).

In the PS approach, we first choose a sequence of increasing radii, 0 = *r*_0_ < *r*_1_ < ⋯ < *r_v_* < ⋯ < *r_V_*. We let radius *r*_0_ = 0 and weights *w*(*s*; *s*′; 0) = **I**(*s* = *s*′) initialize the process, where **I**(·) is an indicator function. At the *v*th iteration, we first update the estimator for the level of differential abundance by the weight *w*(*s*; *s*′; *v* – 1)

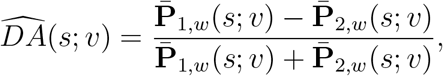

where the weighted mean of relative abundance is estimated by *w*(*s*; *s*′; *v* – 1)

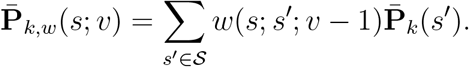

After updating 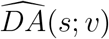, we can then update the weights

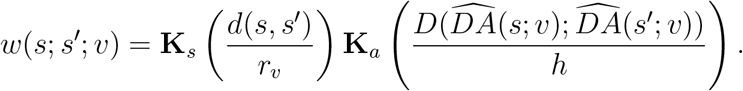

We can update the weights and differential abundance level alternatively.

When does this iterative procedure stop? The stopping criteria for the above iterative procedure are based on the estimated level of differential abundance. Specifically, we choose the *V_L_* as the number of required iterations and *V_U_* as the largest possible iteration number. We shall not stop the PS approach when *v* ≤ *V_L_* and always end it when *v* = *V_U_*. When *V_L_* < *v* ≤ *V_U_*, we compare 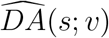 with 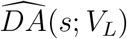 and stop if

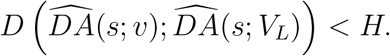

After the PS approach stops, we collect the weights in the final iteration and use them to construct a weighted abundance table.

#### 4.3.3 Weighted RDB test

After ASV aggregation by the weights in Section 4.3.2, we can test the reference-based hypothesis in (2) on the weighted abundance table. To be robust to the zero counts, we adopt the iterative empirical Bayes framework introduced in Wang (2023). We first review the idea of the iterative empirical Bayes framework and then introduce the extension of this idea on the weighted abundance table.

##### Vanilla RDB test with infinite samples

To introduce the iterative empirical Bayes framework, we consider an ideal setting where the number of observed samples is infinite. In other words, we assume that **Q**_1_(*s*) and **Q**_2_(*s*) are observed in advance. The key observation in the iterative empirical Bayes framework is that the fold changes of non-differential abundant ASVs are always the same regardless of renormalization. Putting it mathematically, for any subset 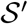 such that 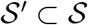, there exists a constant *b*′ such that

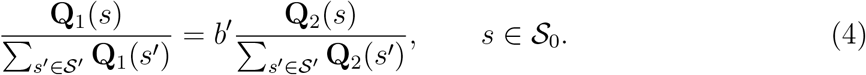

Motivated by this observation, we introduce the vanilla RDB test with infinite samples. In the RDB test, we define the active set 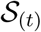 and differential abundant set 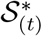 at the *t*th iteration. To initialize the process, we choose 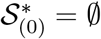 and 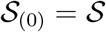. In the *t*th iteration, we renormalize the relative abundance with respect to the active set 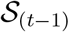

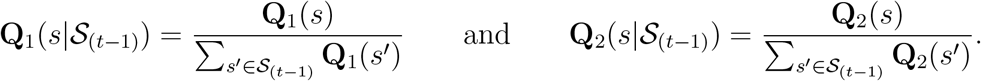

The observation in (4) suggests that 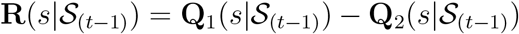 has the same sign for non-differential abundant ASVs. Since 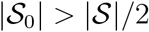 in the definition of the reference set, we can conclude

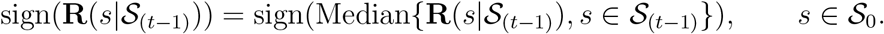

Equivalently, if we define

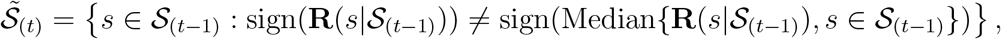

we can conclude ASVs in 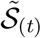 are differential abundant ASVs. Then we can update the active set and differential abundant set

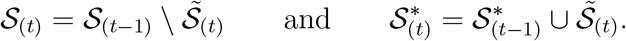

We can repeat this procedure and stop when 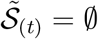. If the last iteration is *T*, the estimator for 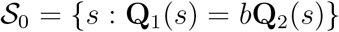 and 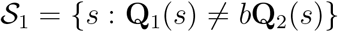 are 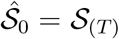 and 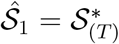 When we observe infinite samples, Wang (2023) proves that the RDB test can identify all differential abundant ASVs accurately, i.e., 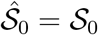 and 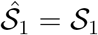.

##### Vanilla RDB test with finite samples

In practice, we only observe finite samples and need to redefine the RDB test to account for the randomness in data. Specifically, we can replace the sign comparison with the directional two-sample testing in 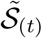. To see the connection between the sign comparison and the directional two-sample testing, we rewrite 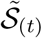 in the following way

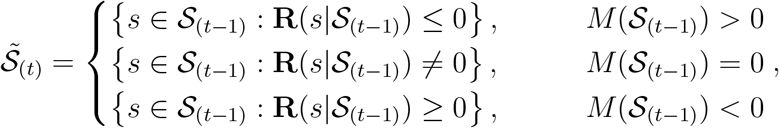

where 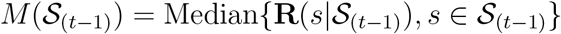. The above form of 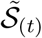 suggests the empirical Bayes interpretation of this method: 1) we first look at the distribution of 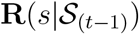 to infer testing direction; 2) we conduct the directional two-sample test at each *s*. Due to the finite samples, we can evaluate *t*-test statistics on the relative abundance instead of 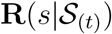

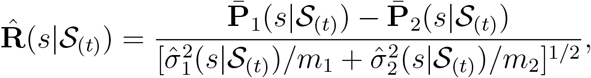

where 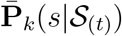 and 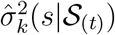 are the mean and variance of renormalized relative abundance. By replacing it with *t*-test statistics, we can redefine 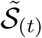 as the result of directional two-sample testing

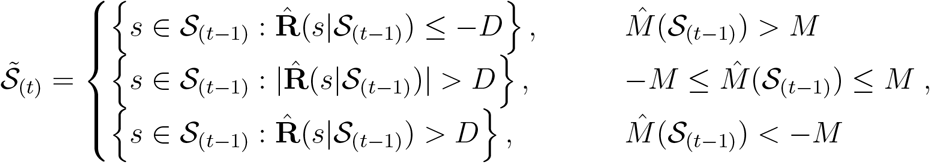

where 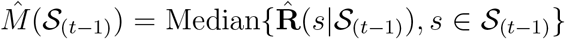. Here *M* > 0 is a threshold for the median, and *D* > 0 is the critical value for the directional two-sample test. Due to the iterative nature, controlling false discovery in the RDB test is slightly different from conventional multiple testing settings (*p*-values are not updated iteratively in conventional settings). We need to choose *M* and *D* to control the false discoveries, and their choices are discussed in Wang (2023). It is worth noting that the design of the RDB test allows some correlation between test statistics since independence structure is not a reasonable assumption for microbiome data due to the negative correlation in the compositional data and some strong dependence between microbial species. Compared with classical differential abundance analysis, the RDB test can simultaneously handle the zero counts and compositionality of data.

##### Weighted RDB test

Since we already assign the weights between ASVs, we consider utilizing information from similar ASVs to determine the sign of 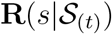. Instead of using the standard two-sample *t*-test, we adopt weighted *t*-test statistics

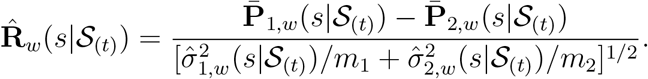

Here, 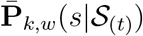 and 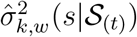 are weighted mean and variance of weighted renormalized relative abundance at ASV *s*, defined as

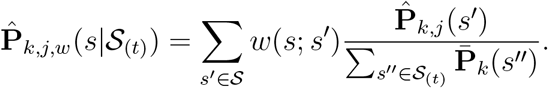

Through borrowing strength from its neighborhood, weighted *t*-test statistics are more powerful than standard *t*-test statistics. Given the weighted *t*-test statistics, we can replace all standard *t*-test statistics with weighted *t*-test statistics in 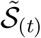

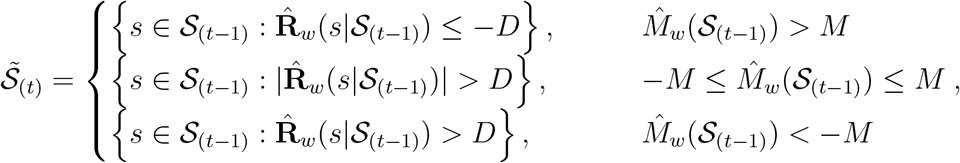

where 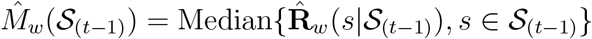. After using weighted *t*-test statistics, we have a new method called the weighted RDB test. In the weighted RDB test, we can still apply the same false discovery control mechanism as the original RDB test since its design allows some correlation between test statistics.

### 4.4 Remarks on MsRDB Test

#### 4.4.1 Results Interpretation in MsRDB Test

To transfer the results obtained by the MsRDB test into knowledge, we still need to interpret these differentially abundant ASVs. We consider two aspects of MsRDB results interpretation: abundance change and taxonomy assignment.

- (**Abundance change**) The framework of the reference-based hypothesis provides a natural way to interpret the results obtained from the compositional data. Specifically, we can regard the collection of non-differential ASVs 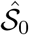 as a reference set and compare everything with this estimated reference set. More specifically, we can always compare the relative abundance with respect to the reference set directly

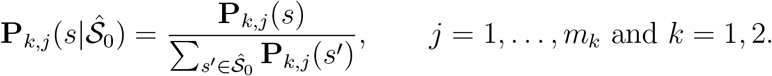 A change in the compositional data is interpreted as a change with respect to the estimated reference set. In the box plots of this work, we always plot the relative abundance with respect to the reference 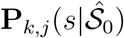 rather than the original relative abundance **P**_*k,j*_(*s*).
- (**Taxonomy assignment**) Since we may find difficulty in assigning taxonomy to some of the ASVs, we opt first to cluster these differentially abundant ASVs into small groups. Specifically, we construct a graph for differentially abundant ASVs by thresholding the ASV distance matrix and then apply a graph-based clustering method for the resulting graph. After the small groups are constructed, we assign taxonomy to each small group of differential ASVs. The benefit of assigning taxonomy to the clusters is that it allows detecting unknown differentially abundant ASVs and assigns different taxonomy rank to different ASV groups.

#### 4.4.2 Covariate Balancing

In an observational study, we also observe several additional covariates **X**_*k,j*_, which might be related to treatment assignment and compositional data. the MsRDB test can work with covariate balancing techniques, such as the weighting method (Imbens and Rubin, 2015), to reduce the potential confounding effect. The weighting method here is different from the ASV-weighted method discussed in the previous section. The weighting method aims to assign weights for each subject so that the distributions of the covariates in each population are roughly the same. Specifically, after the weights *w_k,j_* are assigned to each subject, we can consider a subject-weighted version of mean and variance instead of standard mean and variance in the original MsRDB test. In particular, we choose the empirical balancing calibration weighting method (CAL) proposed by Chan et al. (2016), which provides a nonparametric way to estimate weights.

#### 4.4.3 Practical Considerations

To implement the MsRDB test, we need to specify the choices of several tuning parameters.

- **(multiscale adaptive weights)** We need to choose the kernel **K**_*s*_ and radii *r*_1_ < ⋯ < *r_V_* together. To simplify the implementation, we adopt the idea from the *k*-nearest neighbor algorithm. Specifically, we choose **K**_*s*_(*x*) = **I**(*x* < 1). The radius *r_v_* at ASV *s* is chosen in the following way: we first sort the distance *d*(*s, s′*) in a sense that

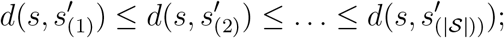

the radius *r_v_* = *d*(*s, s′*_(*k_v_*)_) is chosen for a predefined integer *k_v_*. Equivalently, we include *k_v_* neighbors at the iteration *v* and assign the weights of these *k_v_* neighbors as 1. We choose *k_v_* as a geometric sequence, that is, *k_v_* = ⌈*k^v/V^*⌉, where *V* = *V_U_* is the largest number of iterations, and *k* is the number of neighbors in the last iteration. The main advantage of using a strategy in the *k*-nearest neighbor algorithm is that the total number of weights we need to store is just 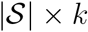, so the computation complexity of the weights update is also 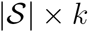. There are many different options for **K**_*a*_ and *h*. Based on our experience, we choose **K**_*a*_(*x*) = exp(–2*x*^2^) and 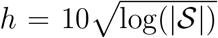 in our implementation. These parameters are mainly used to separate the ASVs with different levels of differential abundance. In the stopping criteria, we choose *V_L_* = 5, 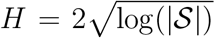 and *V_U_* = *V* =10. *V_L_* cannot be set as a too small number, e.g. 1, since the first a few iterations might not be stable. The computational complexity relies on *V_U_*, as it controls the number of iterations we need to evaluate. See more discussions in Li et al. (2011); Wang et al. (2019).
- **(weighted RDB test)** In the weighted RDB test, we use the same choices of parameters as the original RDB test. As we always aim to control FDR, the critical value for directional testing is chosen adaptively, as discussed in Section 4 of Wang (2023). The covariate balancing method we used in the MsRDB test is the weighting method implemented in `ATE` R package.

## Acknowledgments

S. Wang acknowledges support from NSF Grant DMS-2113458.

## Data Availability

All three data sets can be downloaded from Qiita (https://qiita.ucsd.edu/). Data set in Yatsunenko et al. (2012) is under study ID 850. Data set in Vangay et al. (2018) is under study ID 12080. Data set in Bokulich et al. (2014) is under study ID 2019.

## Code Availability

All analyses can be found under https://github.com/lakerwsl/MsRDB-Manuscript-Code.

## Supplement Material

### Simulation Setup

The simulated ASV data set is generated from a real gut microbiota data set collected in Yatsunenko et al. (2012). This data set includes V4 16S rRNA data from 528 subjects. The data set is trimmed at length 100 and denoised by Deblur method (Amir et al., 2017).

### Simulation Study I

We keep 1965 ASVs which appear in more than 3% of all subjects. To simulate the data set, we consider the following procedure:

1. We randomly draw *m* subjects as the treated group and *m* subjects as the control group.
2. We randomly select *s* ASVs and their neighbors as differential ASVs. The distance between ASVs is defined by the hamming distance between aligned sequences and calculated by function `DistanceMatrix` in `DECIPHER` R package. The number of neighbors is selected randomly between 10 and 15.
3. For each selected differential ASV and its neighbors, the signal strength is randomly chosen between 1 + λ and 1 + 2λ. The count data in the treated group are multiplied by the chosen signal strength.
4. To mimic the experimental bias, we multiply the count data of each ASV by a measurement efficiency constant randomly drawn between 1 and 10.

In this simulation setup, *m* is the sample size, λ is the signal strength, and *s* is the number of differential ASVs (sparsity). Specifically, we consider three sets of simulation experiments:

- We first investigate the influence of sample size, so we choose *m* = 50, 100, 200, 400, λ = 10, and *s* = 20. The results are summarized in Figure 4.
- We next investigate the influence of signal strength, so we choose *m* = 300, λ = 2, 4, 6, 8, and *s* = 20. The results are summarized in Figure S1.
- Finally, we investigate the influence of the number of differential ASVs, so we choose *m* = 200, λ = 8, and *s* = 5, 10, 15, 20. The results are summarized in Figure S2.

**Figure S1:**
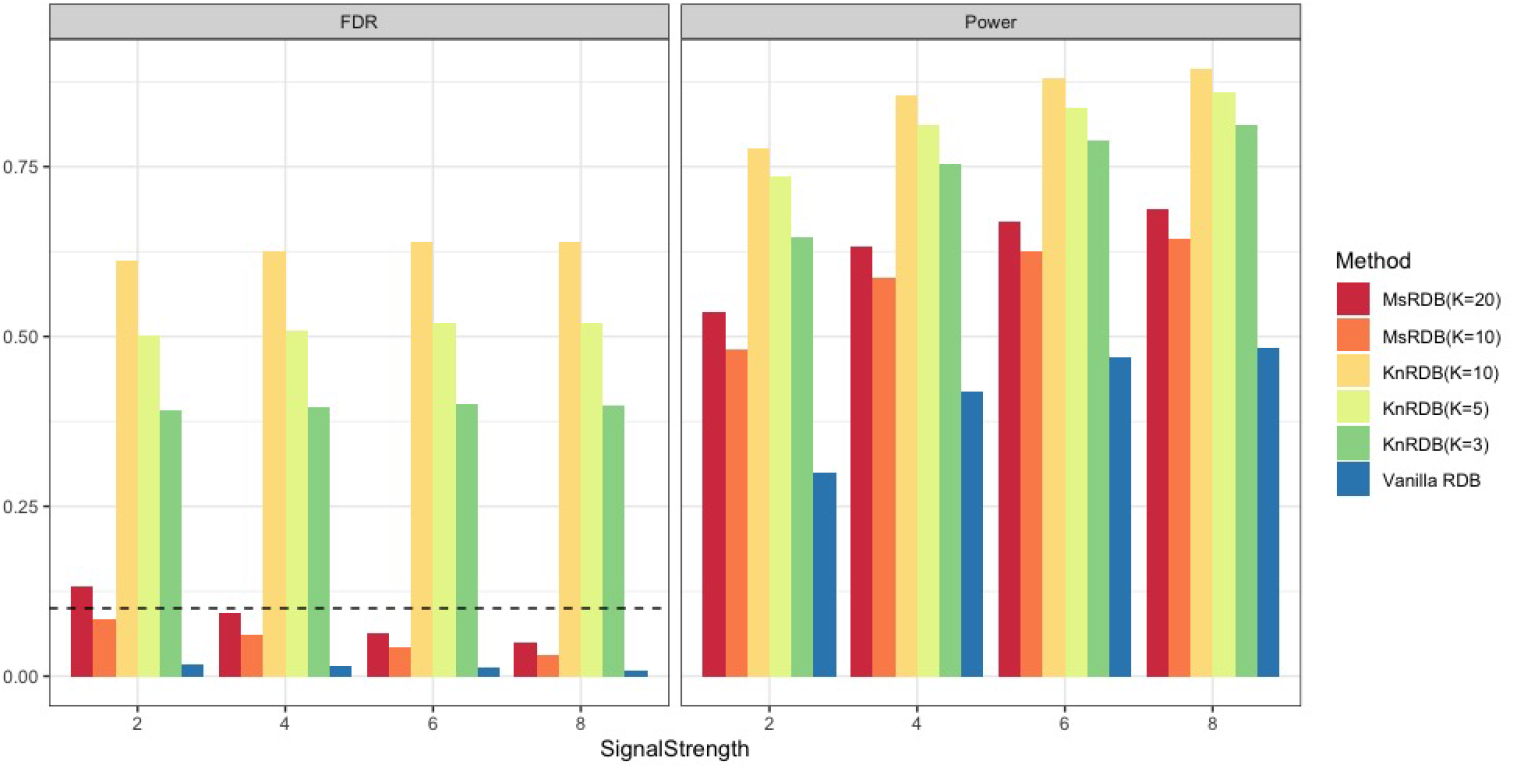
FDR and power comparisons in simulated data set when the signal strength is different. Left and right figures show the FDR and power of various versions of RDB methods when the signal strength are 2, 4, 6, and 8. The dashed line in the left figure is the target level of FDR at 10%. The results in this figure show that the power becomes larger when the signal strength increases.

**Figure S2:**
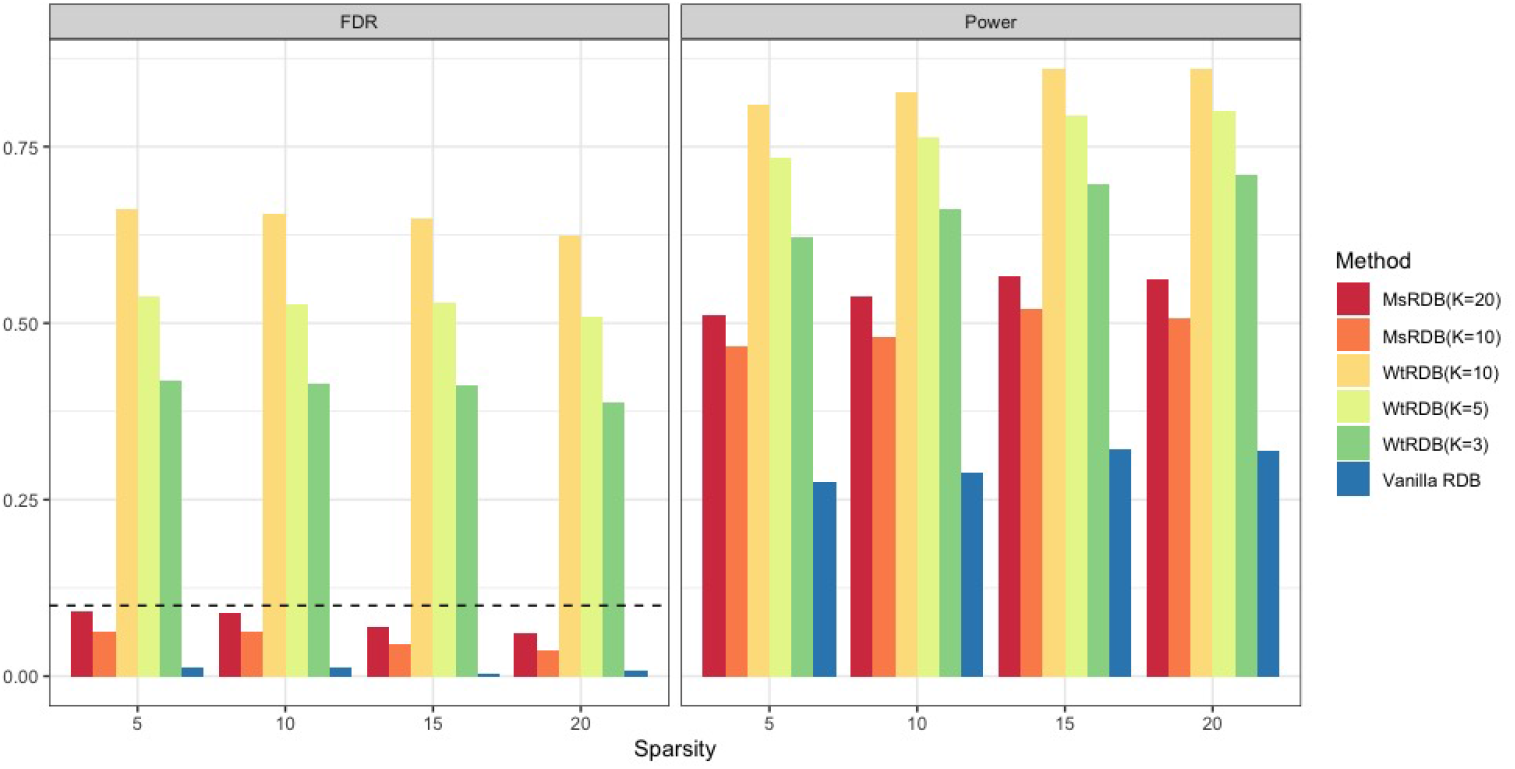
FDR and power comparisons in simulated data set when the number of differentially abundant ASVs is different. Left and right figures show the FDR and power of various versions of RDB methods when the numbers of differentially abundant ASVs are 5, 10, 15, and 20. The dashed line in the left figure is the target level of FDR at 10%. The results in this figure show that the performance of various methods is relatively stable when the number of differentially abundant ASVs is different.

### Simulation Study II

We keep 859 ASVs which appear in more than 10% of all subjects. To simulate the data, we adopt the same procedure as simulation study I with the following modification:

- In Figure 5, the differential ASVs are all ASVs in genera *Ruminococcus*, *Bacteroides*, *Faecalibacterium, Oscillibacter*, and *Blautia*. The signal strength of each genus is randomly chosen between 1 + λ and 1 + 2λ with λ = 20. There is no measurement error.
- In Figure S3, the differential ASVs are all ASVs in classes *Bacteroidia* and *Bacilli*. The signal strength of each class is randomly chosen between 1 + λ and 1 + 2λ with λ = 20. There is no measurement error.
- In Figure S4, the differential ASVs are *s* = 5 ASV clusters chosen as Step 2. The number of neighbors is selected randomly between 5 and 10. The signal strength of each class is randomly chosen between 1 + λ and 1 + 2λ with λ = 20. There is no measurement error.
- In Figure S5, the setting is the same with Figure S4, but there is measurement error as Step 4.
- In Figure S6, the setting is the same with Figure S5, but the sequencing depth is multiply by *η* in the first group, where *η* is randomly chosen as 2 or 3.

In this set of simulation experiment, we consider the following methods and tuning parameters

- MsRDB: *k* = 10 and other tuning parameters are default values.
- RDB: all tuning parameters are default values.
- ANCOM.BC: multiple testing method is “holm” (same with its tutorial) and the pseudo-count is 1. Other tuning parameters are default values.
- DACOMP: the pseudo-count is 1 in reference selection, the test is `DACOMP.TEST.NAME.WILCOXON`, and multiple testing method is “BH”. Other tuning parameters are default values.
- ALDEx2: multiple testing method is “BH” and other tuning parameters are default values.
- StructFDR: the phylogenetic tree is estimated by UPGMA method (implemented by `upgma` in `phangorn` R package) and the distance matrix is calculated by function `DistanceMatrix` in `DECIPHER` R package. Set raw.count as TRUE and other tuning parameters are default values.

### Simulation Study III

In this set of simulation experiment, we follow the same procedure in Figure S4 with *s* = 10 ASV clusters as differential abundant ASVs. The true differential abundant genus (family) is a genus (family) with at least one differential abundant ASV.

We consider the following methods and tuning parameters

- MsRDB: *k* = 10 and other tuning parameters are default values. The method is applied to ASV table directly. All genera (families) including differential abundant ASVs are differential abundant genera (families).
- RDB-ASV: all tuning parameters are default values. The method is applied to ASV table directly. All genera (families) including differential abundant ASVs are differential abundant genera (families).
- RDB-Taxa: all tuning parameters are default values. The ASV table is converted into a genus (family) table and then the method is applied to genus (family) table.
- ANCOMBC-ASV: multiple testing method is “holm” (same with its tutorial) and the pseudo-count is 1. Other tuning parameters are default values. The method is applied to ASV table directly. All genera (families) including differential abundant ASVs are differential abundant genera (families).
- ANCOMBC-Taxa: multiple testing method is “holm” (same with its tutorial) and the pseudo-count is 1. Other tuning parameters are default values. The ASV table is converted into a genus (family) table and then the method is applied to genus (family) table.
- StructFDR: the phylogenetic tree is estimated by UPGMA method (implemented by `upgma` in `phangorn` R package) and the distance matrix is calculated by function `DistanceMatrix` in `DECIPHER` R package. Set raw.count as TRUE and other tuning parameters are default values. The method is applied to ASV table directly. All genera (families) including differential abundant ASVs are differential abundant genera (families).

**Figure S3:**
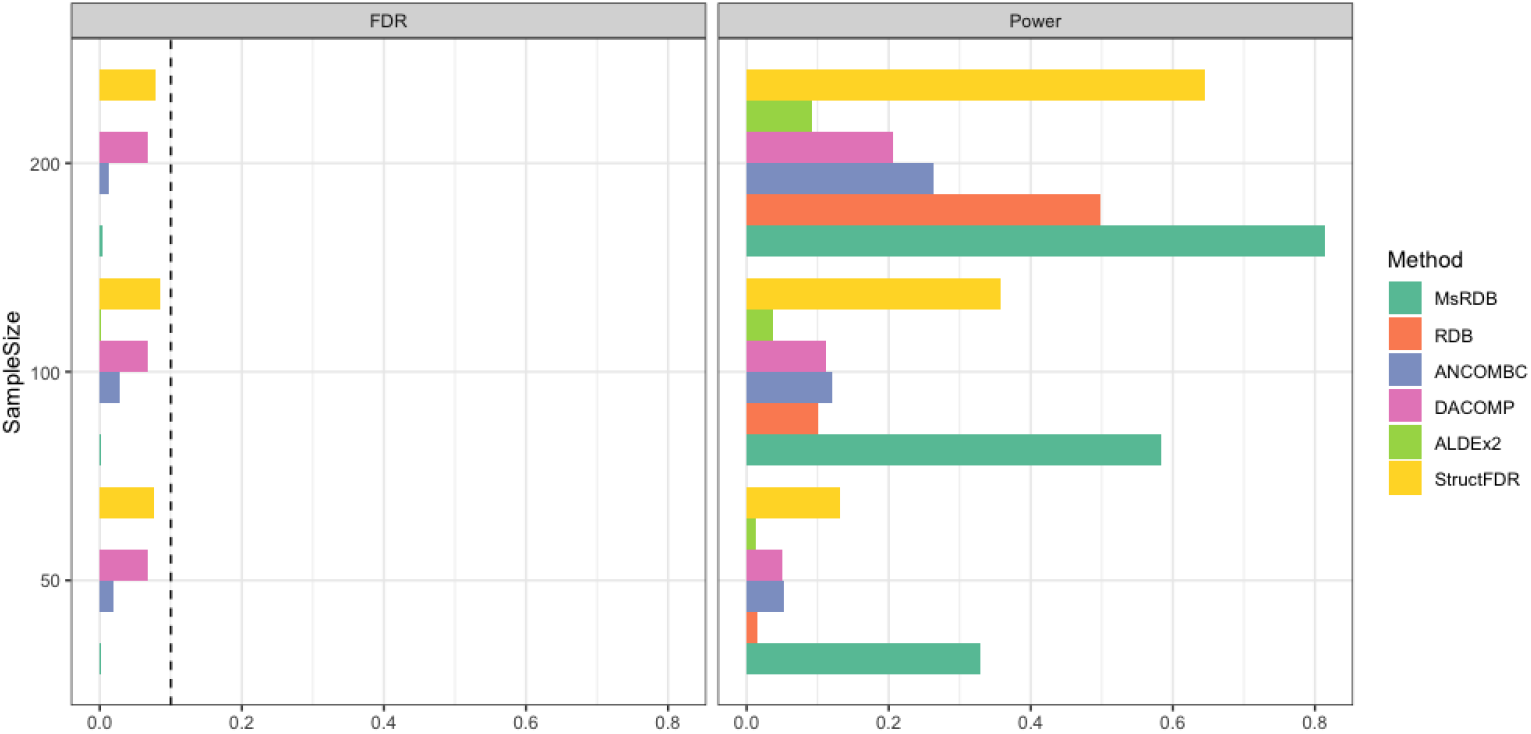
Comparisons of differential abundance tests in simulated data sets when differential abundant microbes are different classes. The differential abundant ASVs are all ASVs in classes *Bacteroidia* and *Bacilli*. Left and right figures show the FDR and power of differential abundance tests when the sample sizes are 50, 100, and 200. The dashed line in the left figure is the target level of FDR at 10%. All methods can control FDR at the ASV level very well. The results confirm that aggregating information from the neighborhood can lead to a more powerful test.

**Figure S4:**
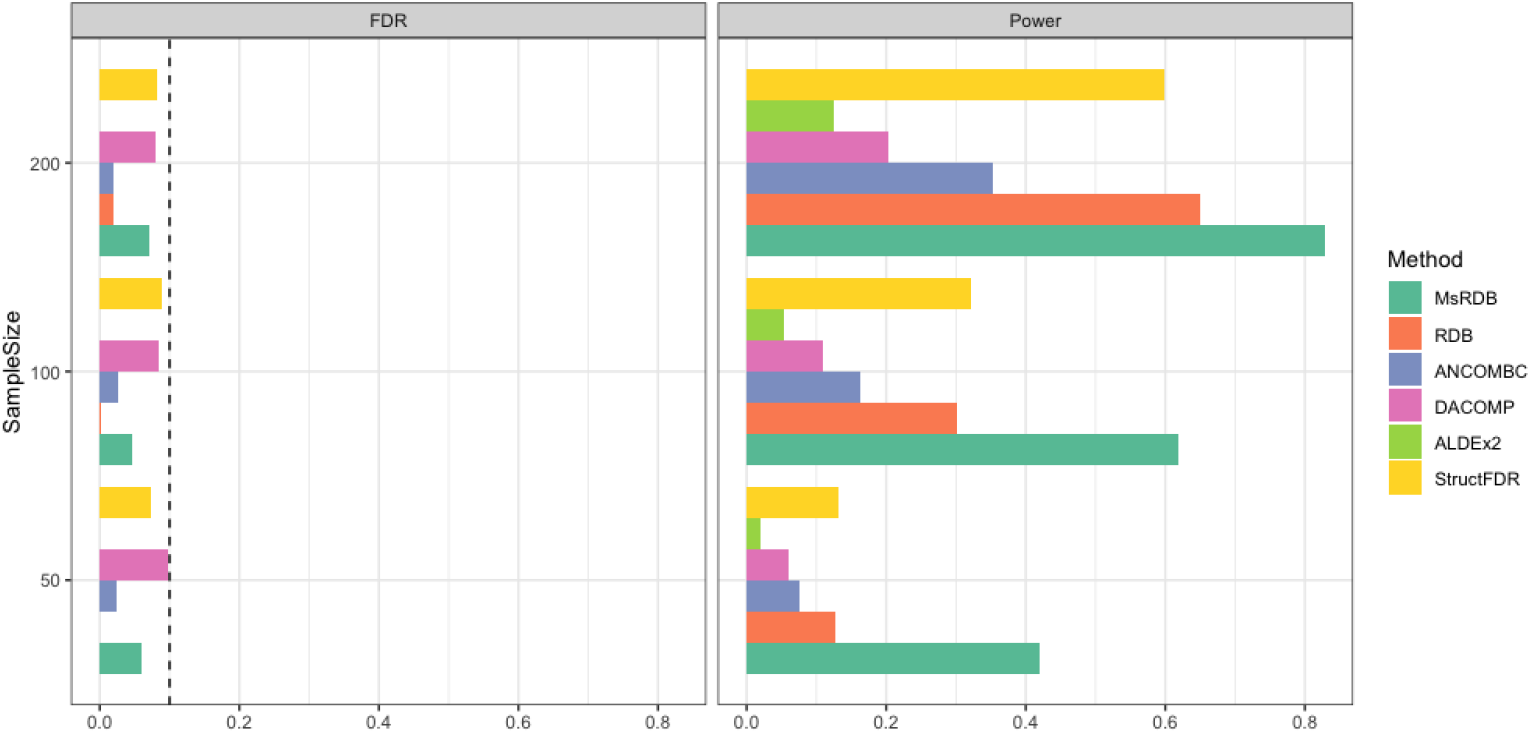
Comparisons of differential abundance tests in simulated data set when differential abundant microbes are different ASV clusters. The differential abundant ASVs are ASVs in small ASV clusters. Left and right figures show the FDR and power of differential abundance tests when the sample sizes are 50, 100, and 200. The dashed line in the left figure is the target level of FDR at 10%. All methods can control FDR at the ASV level very well. The results confirm that aggregating information from the neighborhood can lead to a more powerful test.

**Figure S5:**
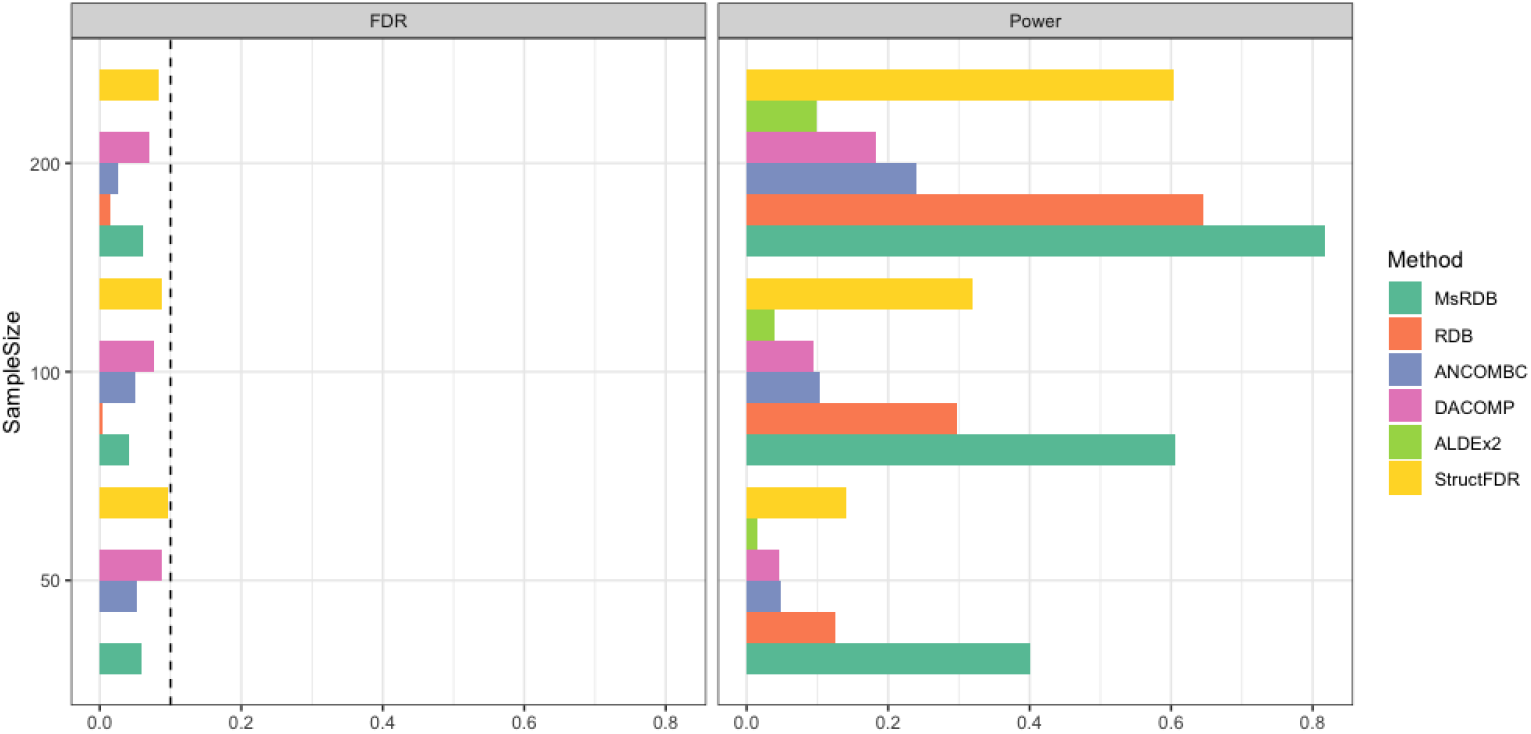
Comparisons of differential abundance tests in simulated data set when the ASV data has measurement error. The differential abundant ASVs are ASVs in small ASV clusters, and the ASV data has measurement error. Left and right figures show the FDR and power of differential abundance tests when the sample sizes are 50, 100, and 200. The dashed line in the left figure is the target level of FDR at 10%. All methods are robust to the measurement error.

**Figure S6:**
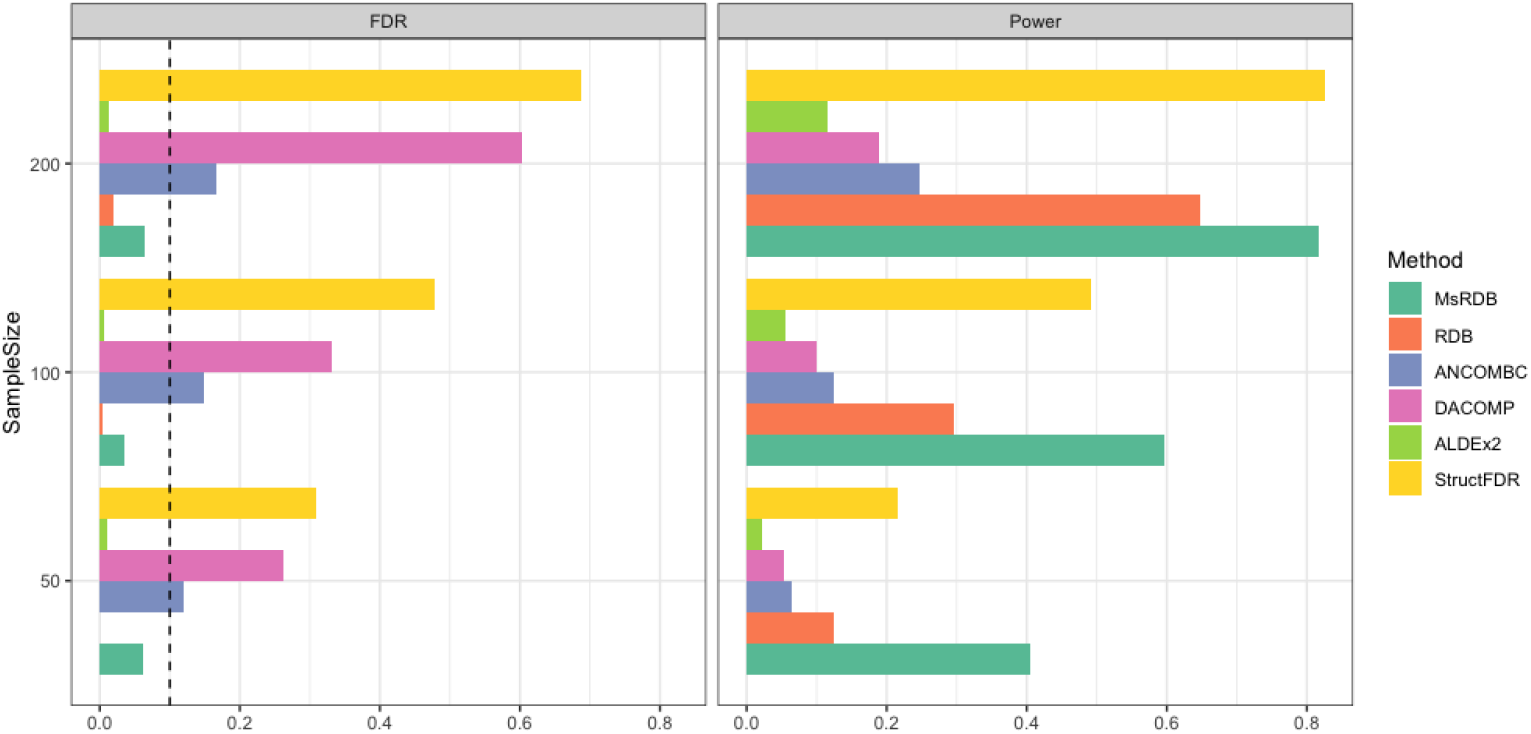
Comparisons of differential abundance tests in simulated data set when the ASV data has measurement error and unbalanced sequencing depth. The differential abundant ASVs are ASVs in small ASV clusters, and the ASV data has measurement error and unbalanced sequencing depth. Left and right figures show the FDR and power of differential abundance tests when the sample sizes are 50, 100, and 200. The dashed line in the left figure is the target level of FDR at 10%. Most methods are robust to measurement error and unbalanced sequencing depth. FDR is inflated in DACOMP and StructFDR due to unbalanced sequencing depth.

**Figure S7:**
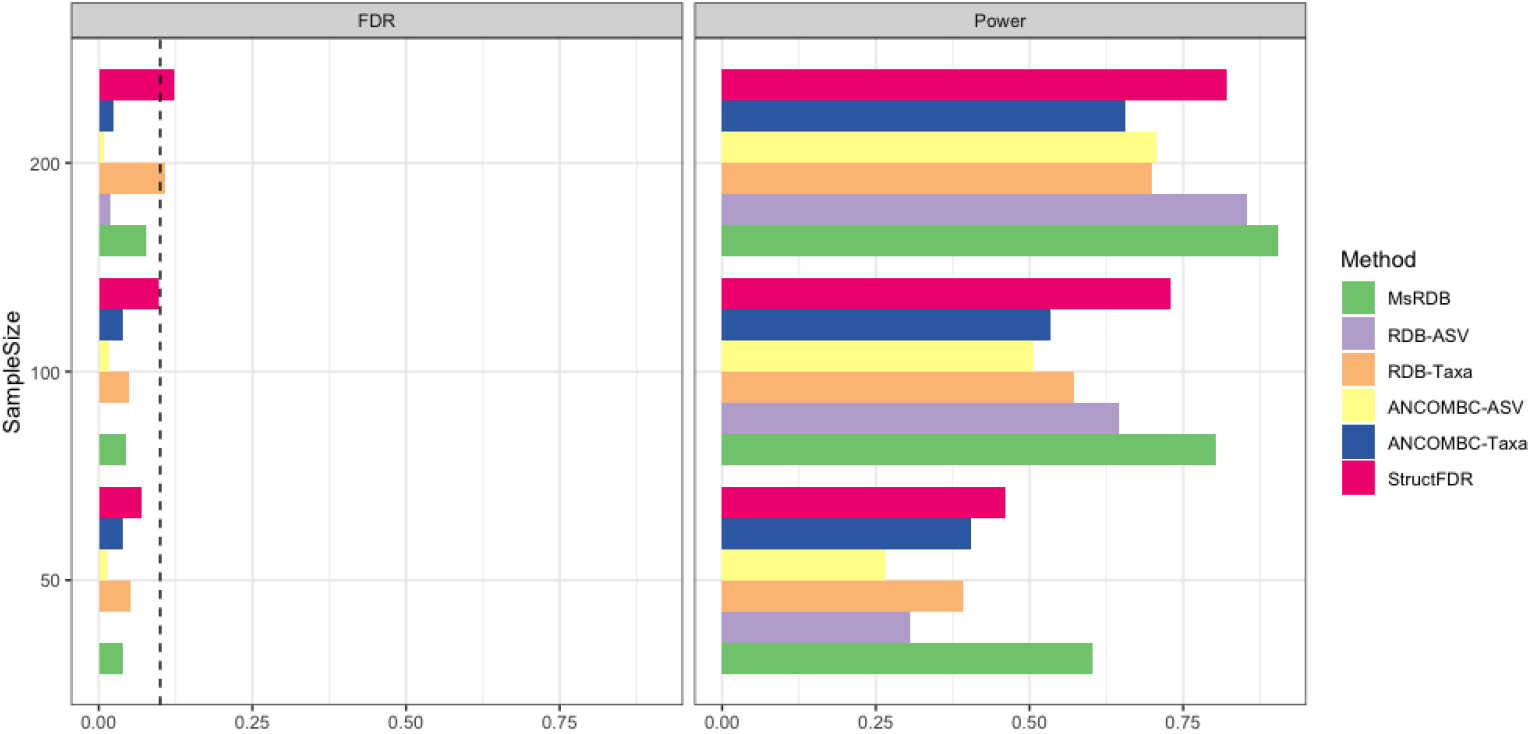
Comparisons of differential abundance tests in identifying differential abundant families. The differential abundant family is a family with at least one differential abundant ASV. Left and right figures show the FDR and power of differential abundance tests when the sample sizes are 50, 100, and 200. The dashed line in the left figure is the target level of FDR at 10%. All methods can control FDR at the family level very well. Choosing family as analysis units can improve the power of RDB and ANCOM.BC when the sample size is small.

### Differential Abundance Analysis in the Study of Immigration

Taxonomy in the immigration study data set (Vangay et al., 2018) is assigned by function `assignTaxonomy` in `dada2` R package with training files from the Silva Project’s version 138.1 release. The differential abundance analysis includes 14471 ASVs assigned to kingdom *Bacteria*. Besides the microbiome data, we also observe several extra variables in this observational study. In particular, we include age and BMI into our analysis, as they might bring potential confounding effects. The degree of covariance balancing in age and BMI suggests that neither variable is well balanced, so we must adjust these two covariates in differential abundance analysis (Figure S8). We adopt the weighting method implemented in `ATE` R package for the RDB and the MsRDB tests to balance covariates. In the MsRDB test, the distance matrix between ASVs’ sequences is calculated by function `DistanceMatrix` in `DECIPHER` R package. In the MsRDB test, the number of neighbors in the last iteration is *k* = 20.

**Figure S8:**
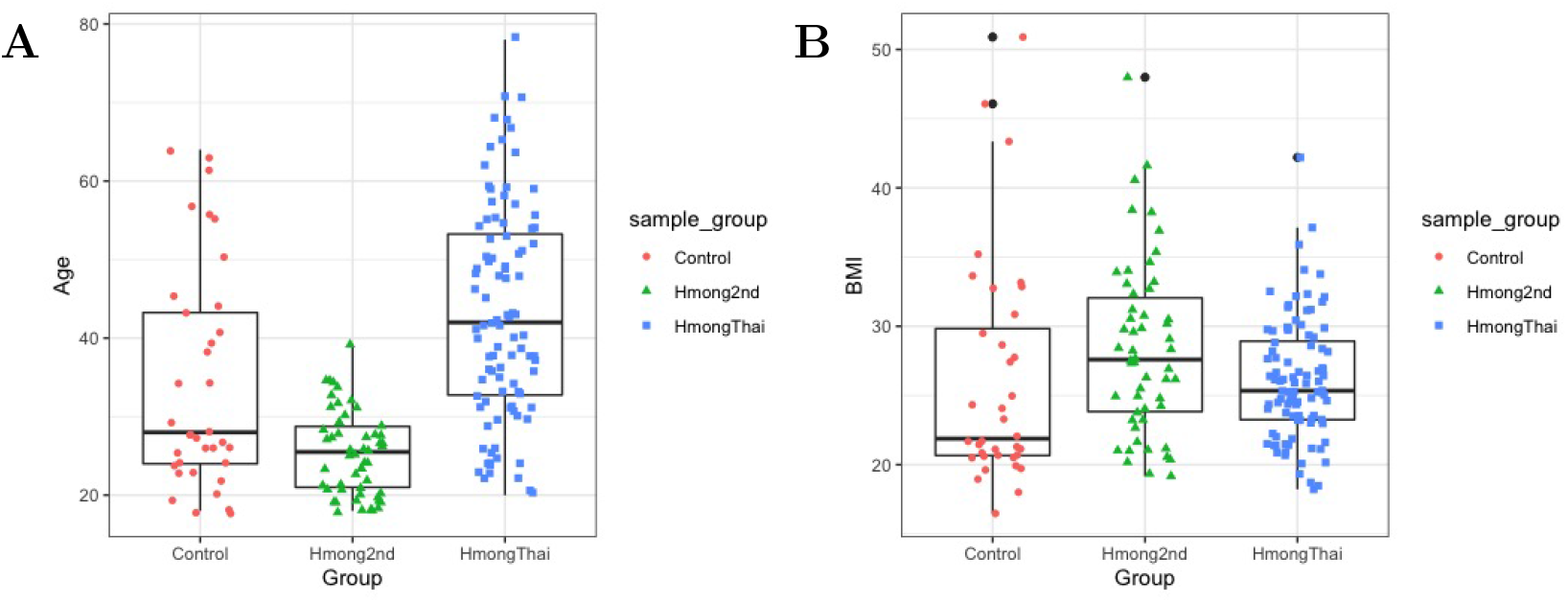
Degree of covariance balancing in age and BMI in immigration study. This figure compares the distribution of covariates among HmongThai, Hmong2nd, and Control. Left is age, and right is BMI.

**Figure S9:**
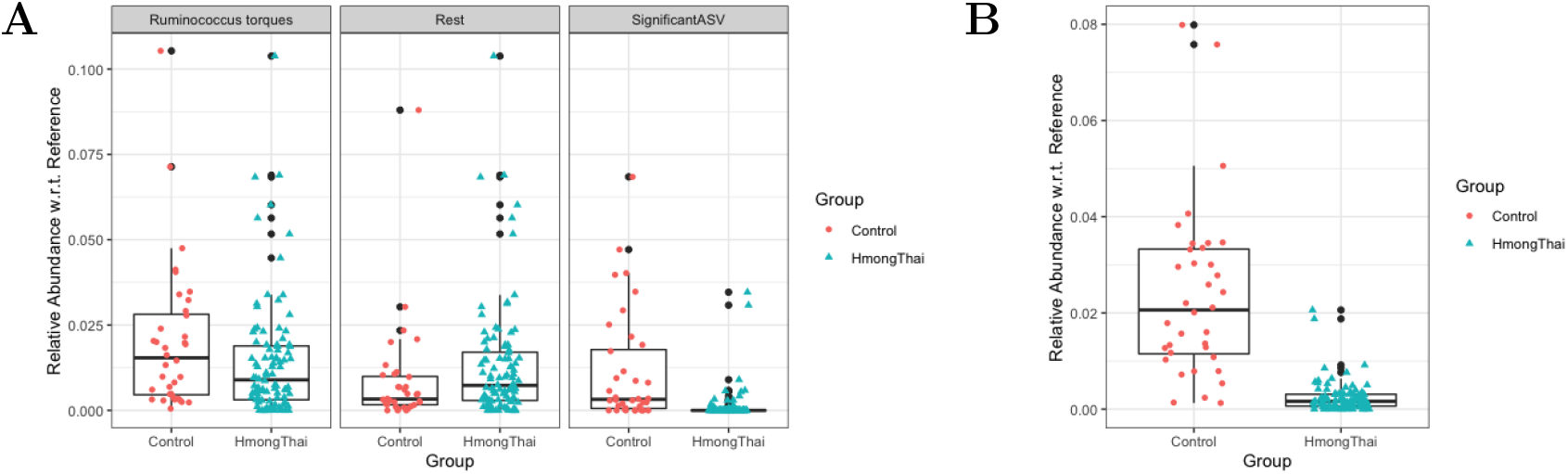
Taxa detected by MsRDB test on the microbiota data set in immigration study. The reference set in all box plots is all non-differentially abundant ASVs revealed by the MsRDB test. (A) The box plots show the relative abundance with respect to the reference set in all ASVs assigned to genus *Ruminococcus torques* (left), in significant ASVs within *Ruminococcus torques* (right), and in the rest of ASVs within *Ruminococcus torques* (middle). (B) The box plot shows the relative abundance with respect to the reference set in all significant ASVs assigned to the NA genus.

**Figure S10:**
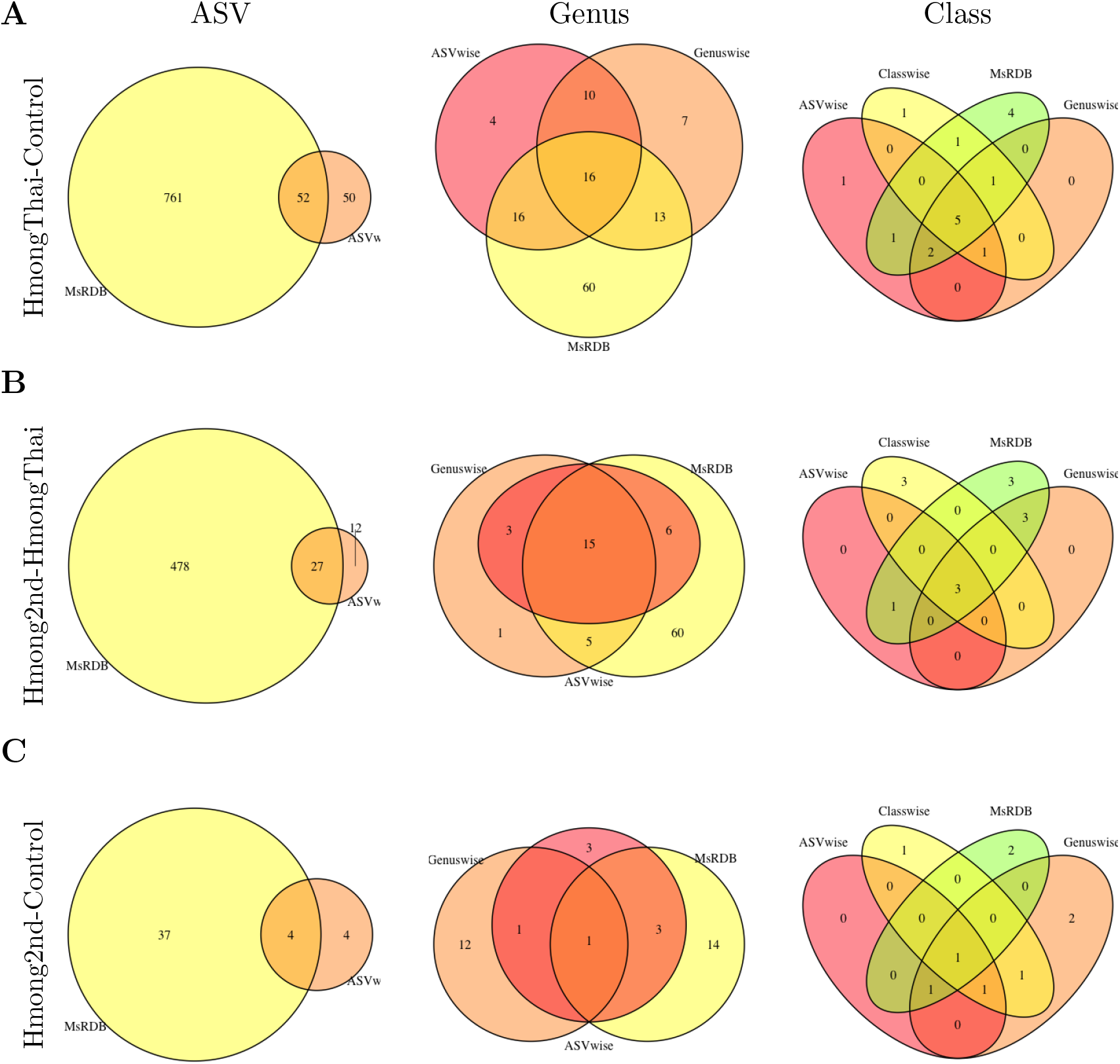
Comparisons between ASV-wise analysis (ANCOM-BC), taxon-wise analysis (ANCOM-BC), and the MsRDB test on the immigration microbiota data set. In this figure, ANCOM-BC is applied in ASV-wise analysis and taxon-wise analysis. (A-C) shows the overlap of results obtained by different analysis strategies in pairwise comparisons of three groups of people. The three rows are the comparisons of HmongThai vs. Control, Hmong2nd vs. HmongThai, and Hmong2nd vs. Control.

### Differential Abundance Analysis in the Study of Wine Grape

Similar to the previous data set, we assign taxonomy of ASVs in wine grape data set (Bokulich et al., 2014) by function `assignTaxonomy` in `dada2` R package. The distance matrix in the MsRDB test is also calculated by function `DistanceMatrix` in `DECIPHER` R package. In the MsRDB test, the number of neighbors in the last iteration is *k* = 15. In clustering analysis, we first construct a graph where two significant ASVs are connected if they are in each other *k* nearest neighbor. Then, each connected subgraph is a cluster, and the small clusters with a total number of reads smaller than 10 are removed from the results.

**Figure S11:**
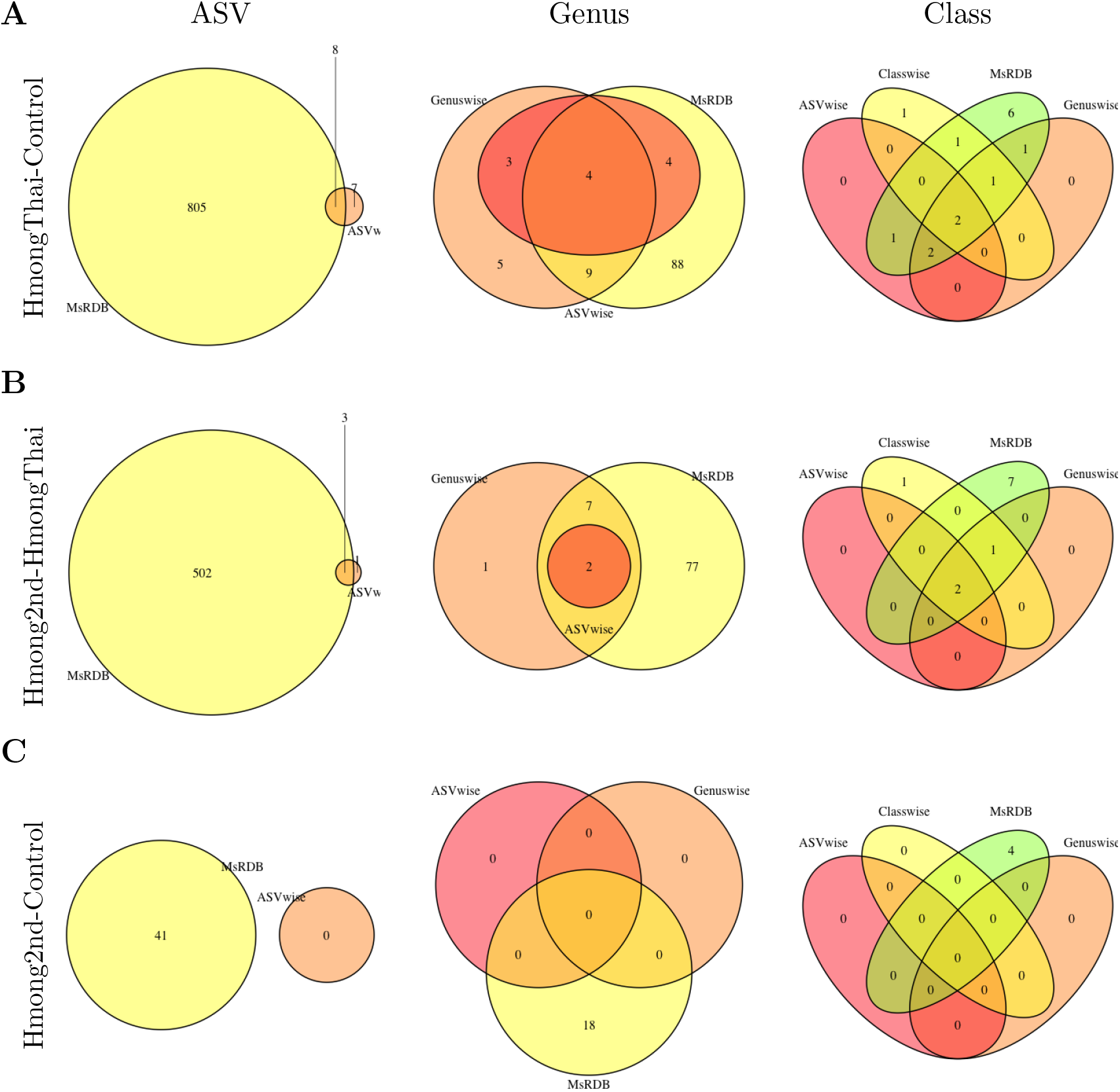
Comparisons between ASV-wise analysis (ALDEx2), taxon-wise analysis (ALDEx2), and the MsRDB test on the immigration microbiota data set. In this figure, ALDEx2 is applied in ASV-wise analysis and taxon-wise analysis. (A-C) shows the overlap of results obtained by different analysis strategies in pairwise comparisons of three groups of people. The three rows are the comparisons of HmongThai vs. Control, Hmong2nd vs. HmongThai, and Hmong2nd vs. Control.

**Figure S12:**
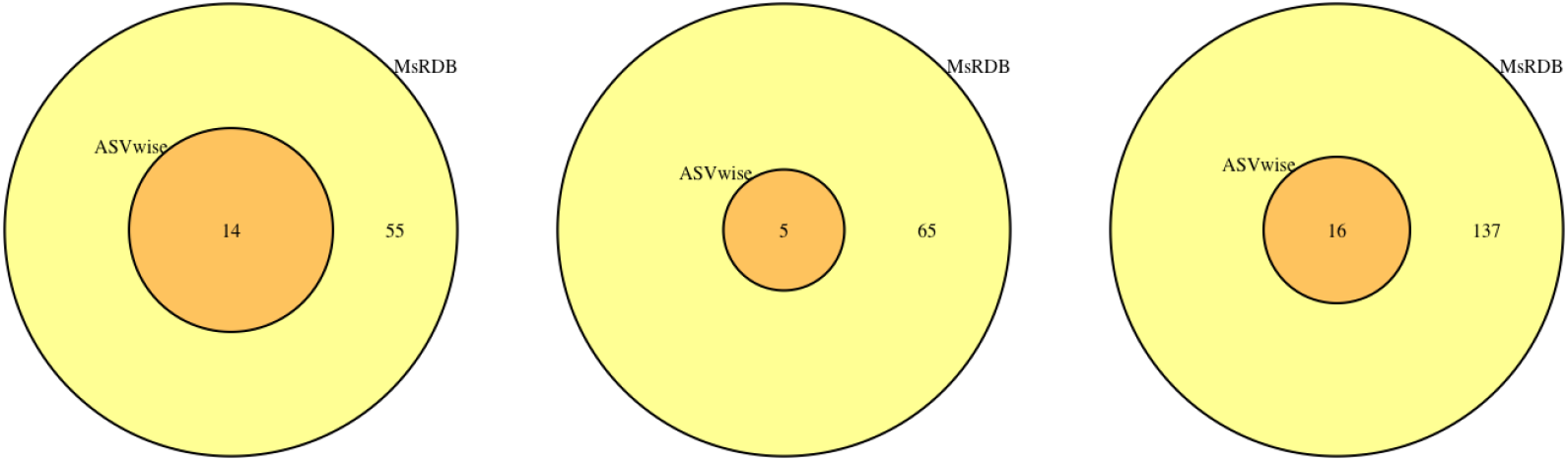
Comparisons between ASV-wise analysis and MsRDB test on the grape microbiota data set. This figure shows the overlap of identified ASV by ASV-wise analysis and the MsRDB test. Left is Sonoma vs. Napa, middle is Central Coast vs. Napa, and right is Central Coast vs. Sonoma.

